# LuxT is a Global Regulator of Low-Cell Density Behaviors Including Type III Secretion, Siderophore Production, and Aerolysin Production in *Vibrio harveyi*

**DOI:** 10.1101/2021.11.09.467953

**Authors:** Michaela J. Eickhoff, Chenyi Fei, Jian-Ping Cong, Bonnie L. Bassler

## Abstract

Quorum sensing (QS) is a chemical communication process in which bacteria produce, release, and detect extracellular signaling molecules called autoinducers. Via combined transcriptional and post-transcriptional regulatory mechanisms, QS allows bacteria to collectively alter gene expression on a population-wide scale. Recently, the TetR-family transcriptional regulator LuxT was shown to control *V. harveyi qrr*1, encoding the Qrr1 small RNA that functions at the core of the QS regulatory cascade. Here, we use RNA-Sequencing to reveal that, beyond control of *qrr*1, LuxT is a global regulator of 414 *V. harveyi* genes including those involved in type III secretion, siderophore production, and aerolysin toxin biosynthesis. Importantly, LuxT directly represses *swrZ,* encoding a GntR-family transcriptional regulator, and LuxT control of type III secretion, siderophore, and aerolysin genes occurs by two mechanisms, one that is SwrZ-dependent and one that is SwrZ-independent. All of these target genes specify QS-controlled behaviors that are enacted when *V. harveyi* is at low cell density. Thus, LuxT and SwrZ function in parallel with QS to drive particular low cell density behaviors. Phylogenetic analyses reveal that *luxT* is highly conserved among *Vibrionaceae,* but *swrZ* is less well conserved. In a test case, we find that in *Aliivibrio fischeri,* LuxT also represses *swrZ*. SwrZ is a repressor of *A. fischeri* siderophore production genes. Thus, LuxT repression of *swrZ* drives activation of *A. fischeri* siderophore gene expression. Our results indicate that LuxT is a major regulator among *Vibrionaceae,* and, in the species that also possess *swrZ*, LuxT functions with SwrZ to control gene expression.

**Importance:** Bacteria precisely tune gene expression patterns to successfully react to changes that occur in the environment. Defining the mechanisms that enable bacteria to thrive in diverse and fluctuating habitats, including in host organisms, is crucial for a deep understanding of the microbial world and also for development of effective applications to promote or to combat particular bacteria. In this study, we show that a regulator called LuxT controls over 400 genes in the marine bacterium *Vibrio harveyi* and, moreover, that LuxT is highly conserved among *Vibrionaceae* species, ubiquitous marine bacteria that often cause disease. We characterize the mechanisms by which LuxT controls genes involved in virulence and nutrient acquisition. We show that LuxT functions in parallel with a set of regulators of the bacterial cell-to-cell communication process called quorum sensing to promote *V. harveyi* behaviors at low cell density.

## Introduction

Bacteria have the remarkable ability to rapidly detect and adapt to environmental fluctuations. Often, bacteria employ transcriptional and post-transcriptional regulatory mechanisms to tune gene expression patterns that enhance survival under varying conditions (1, 2). Such combined regulatory mechanisms are used in a process called quorum sensing (QS) to monitor and react to changes in the cell density and the species composition of the vicinal community. QS involves the production, release, accumulation, and group-wide detection of signaling molecules called autoinducers (AIs). QS fosters the synchronous execution of collective behaviors, typically, ones that are unproductive for an individual bacterium to carry out alone but that become effective when enacted by the group, e.g., bioluminescence, virulence factor production, and biofilm formation (3, 4).

*Vibrio harveyi*, the focus of the current work, is a model QS bacterium that uses three AIs that act in parallel to control bioluminescence, type III secretion, siderophore production, and hundreds of other traits (5–8). Each AI is detected by a cognate membrane-bound receptor. At low-cell density (LCD), AI concentrations are low, and the three unliganded QS receptors act as kinases that drive phosphorylation of the response regulator LuxO. LuxO-P activates transcription of genes encoding five small regulatory RNAs (sRNAs) called the Qrr sRNAs that function post-transcriptionally to activate and repress, respectively, the translation of the master QS regulators AphA and LuxR (9–12). Thus, at LCD, AphA is made and LuxR is not. AphA is responsible for executing the LCD QS program (Fig S1A). At LCD, the Qrr sRNAs also directly control 16 other target mRNAs, and they operate as feedback regulators within the QS signaling pathway (13–15). At high-cell density (HCD), accumulated AIs bind to their cognate QS receptors. In the liganded state, the QS receptors acts as phosphatases, and phosphate is stripped from LuxO, which inactivates it (16, 17). Thus, at HCD, Qrr sRNA production terminates, AphA is not made, and, by contrast, LuxR is produced (8, 10, 12). LuxR drives the HCD QS regulon (Fig S1B).

The five *V. harveyi* Qrr sRNAs possess high sequence identity, and Qrr2-5 regulate an identical set of mRNA targets (10, 13). Qrr1 is the outlier. Because it lacks nine nucleotides that are present in Qrr2-5, Qrr1 fails to regulate *aphA* and two additional mRNA targets that are controlled by Qrr2-5 (10, 11, 13). The genes encoding the Qrr sRNAs differ in their LCD expression levels: *qrr*4 is the most highly transcribed of the set, whereas only low-level transcription of *qrr*1 and *qrr*5 occurs (10). Recently, we discovered that a transcription factor called LuxT represses expression of *qrr*1. LuxT does not regulate *qrr*2-5. Thus, the exclusive involvement of LuxT in *qrr*1 regulation provides a mechanism enabling Qrr1 to uniquely control downstream targets (18). Indeed, via repression of *qrr*1, LuxT indirectly controls the translation of Qrr1 target mRNAs including those encoding a secreted protease, an aerolysin toxin, a chitin deacetylase, and a component involved in capsule polysaccharide secretion. LuxT also activates transcription of this same set of genes by a Qrr1-independent mechanism (18). LuxT repression of *qrr*1 does not significantly alter LuxR translation, and as mentioned, Qrr1 does not regulate AphA (11, 18). These findings indicate that LuxT repression of *qrr*1 tunes the expression of select Qrr1-controlled mRNAs without altering the entire QS regulon.

To date, *qrr*1 is the only gene identified to be directly controlled by LuxT (18, 19). However, LuxT has been linked to the regulation of additional phenotypes in *V. harveyi* and other *Vibrionaceae* including bioluminescence, siderophore production, virulence factor production, and motility (18, 20–23). These findings, together with our initial knowledge that LuxT can regulate transcription of target genes independently of Qrr1 in *V. harveyi* (18), inspired us to investigate the role of LuxT beyond its control of *qrr*1. Here, we use RNA-Sequencing (RNA-Seq) to identify the LuxT regulon, revealing LuxT to be a global regulator of ∼414 genes in *V. harveyi*. We find that LuxT directly represses the *V. harveyi swrZ* gene encoding a GntR-family transcriptional regulator. We use genetic and molecular analyses to show that, in *V. harveyi*, LuxT activates genes required for type III secretion, siderophore production, aerolysin toxin production, and activation occurs by two mechanisms. One mechanism depends on SwrZ: LuxT represses *swrZ* and SwrZ represses target gene expression. The second mechanism is SwrZ independent. Finally, the LuxT-controlled traits identified here are also QS-controlled and are enacted primarily at LCD. Therefore, LuxT functions in parallel with QS to establish *V. harveyi* LCD behaviors. Finally, we analyze *luxT* and *swrZ* conservation among *Vibrionaceae* bacteria, and we demonstrate that, via *swrZ* repression, LuxT also activates genes required for siderophore production in *Aliivibrio fischeri*.

## Results

### LuxT is a global regulator that directly represses *swrZ* transcription

To define the set of genes regulated by LuxT, we used RNA-Seq to compare the transcriptomes of WT and Δ*luxT V. harveyi*. We considered transcripts with 2-fold or higher changes in expression (*p* < 0.01) to be LuxT-regulated, revealing a total of 414 genes: 243 activated and 171 repressed (Fig 1A and Data Set S1). One gene, *swrZ* (*VIBHAR_RS03920*), stood out due to its dramatic repression by LuxT. *swrZ* was previously shown to be repressed by the LuxT homolog SwrT in *Vibrio parahaemolyticus* (22). Specifically, SwrT repressed transcription of *swrZ*, and SwrZ repressed the *laf* genes encoding the lateral flagellar machinery required for *V. parahaemolyticus* swarming locomotion. Thus, through this cascade – repression of a repressor – SwrT activates *V. parahaemolyticus* swarming (22). While *V. harveyi* encodes *laf* genes, their expression is extremely low, and *V. harveyi* swarming has not been documented (24). The *V. harveyi laf* genes were not revealed by the RNA-Seq to be LuxT regulated. Nonetheless, our data demonstrate that the regulatory arrangement in which LuxT represses *swrZ* exists in *V. harveyi*: qRT-PCR verified that LuxT represses *swrZ* expression by ∼100-fold (Fig 1B).

**Fig 1.**
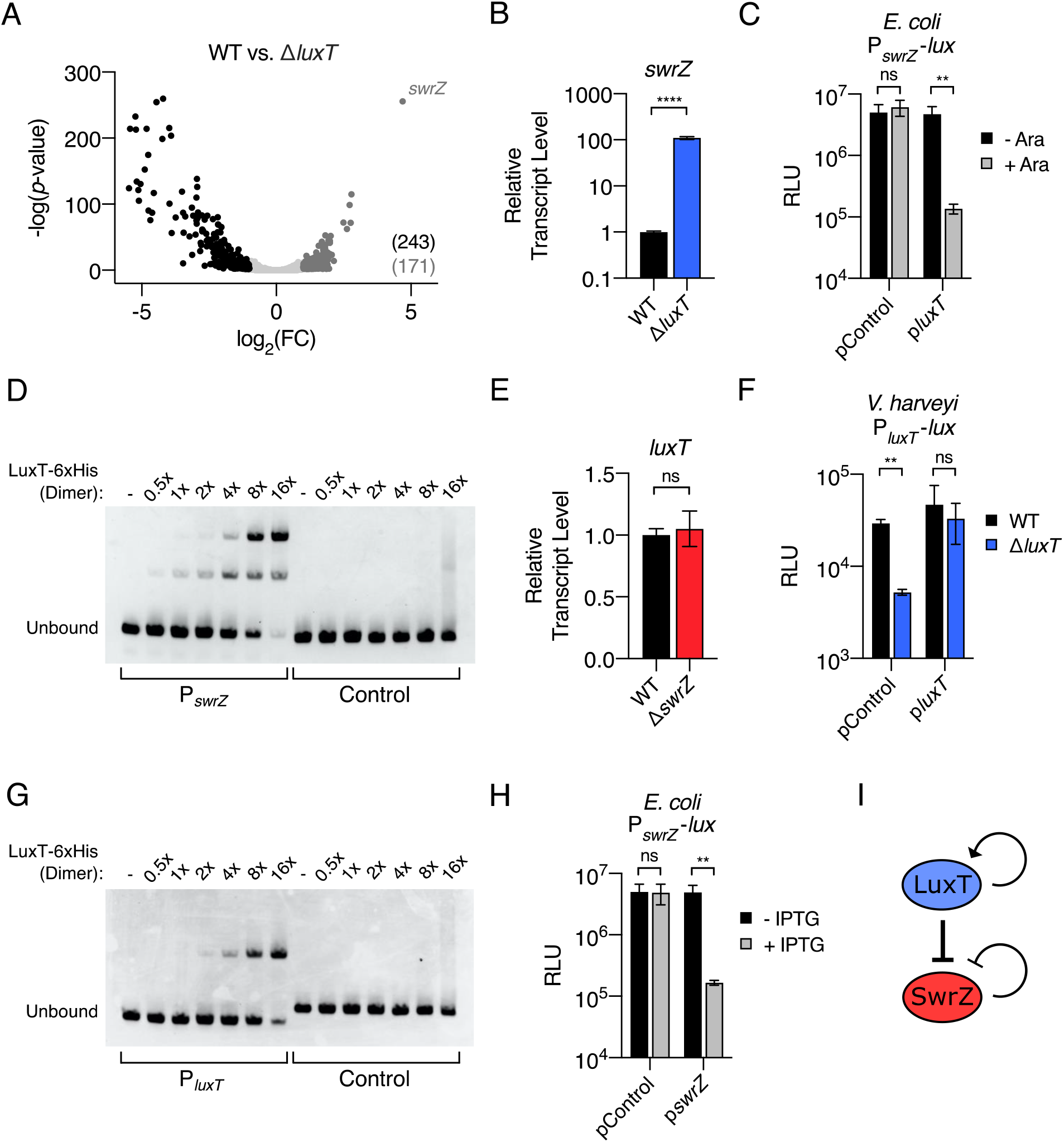
LuxT is a global transcriptional regulator in *V. harveyi* that directly represses *swrZ.* **(A)** Volcano plot depicting RNA-Seq data comparing the transcriptomes of WT and Δ*luxT V. harveyi*. Each data point represents one *V. harveyi* gene. FC designates fold- change. 243 genes were significantly activated (black) and 171 genes were significantly repressed by LuxT (dark gray). **(B)** qRT-PCR measurements of *swrZ* transcript levels in WT (black) and Δ*luxT* (blue) *V. harveyi.* RNA was isolated from strains grown in LM to OD_600_ = 0.1 **(C)** Activity of a plasmid-borne P*_swrZ_-lux* transcriptional reporter in recombinant *E. coli.* pControl denotes that the second introduced plasmid is the empty parent vector and p*luxT* designates that the second vector harbors arabinose-inducible *luxT.* Strains were grown in LB to OD_600_ = 1 in the absence (black) or presence (gray) of 0.01% arabinose. **(D)** EMSA showing binding of LuxT-6xHis to a 113 bp DNA fragment containing the *swrZ* promoter (left) and a 107 bp control fragment containing the *V. harveyi luxC* promoter (right). Previously, it was shown that LuxT does not bind the *luxC* promoter (18). 20 nM DNA probe was incubated with the indicated relative concentrations of the LuxT-6xHis dimer: - indicates no protein, 1x indicates 20 nM, and 16x indicates 320 nM. **(E)** qRT-PCR measurements of *luxT* transcript levels in WT (black) and Δ*swrZ* (red) *V. harveyi.* Growth conditions as in B. **(F)** Activity of a plasmid-borne P*_luxT_-lux* transcriptional reporter in Δ*luxA* (black) and Δ*luxA* Δ*luxT* (blue) *V. harveyi* strains. pControl and p*luxT* denote the empty parent vector and a vector harboring IPTG-inducible *luxT,* respectively. Strains were grown in LM supplemented with 0.5 mM IPTG to OD_600_ = 1. **(G)** As in D, for a 99 bp DNA fragment containing the *luxT* promoter (left) and a 107 bp control fragment containing the *V. harveyi luxC* promoter (right). **(H)** Activity of a plasmid-borne P*_swrZ_-lux* transcriptional reporter in recombinant *E. coli.* pControl denotes that the second introduced plasmid is the empty parent vector and p*swrZ* designates that the second vector harbors IPTG-inducible *swrZ.* Strains were grown in LB to OD_600_ = 1 in the absence (black) or presence (gray) of 0.5 mM IPTG. **(I)** Model for the LuxT/SwrZ regulatory circuit. In C, F, and H, RLU denotes relative light units. In B, C, E, F, and H, error bars represent standard deviations of the means of *n* = 3 biological replicates. Unpaired two-tailed *t* tests with Welch’s correction were performed comparing the indicated conditions. *p*-values: ns ≥ 0.05, ** < 0.01, **** < 0.0001.

To determine if LuxT repression of *swrZ* is direct, we used recombinant *Escherichia coli* to isolate LuxT and the *swrZ* promoter from other *V. harveyi* regulatory components. Two plasmids were introduced into *E. coli.* One plasmid harbored a P*_swrZ_- lux* transcriptional reporter, and the second plasmid carried arabinose-inducible *luxT*. Induction of *luxT* expression drove repression of P*_swrZ_-lux* in the *E. coli* strain carrying the p*luxT* vector, whereas no repression occurred in *E. coli* carrying the empty control vector (Fig 1C). We conclude that LuxT directly represses *swrZ*. Consistent with this supposition, purified LuxT-6xHis protein bound to the *swrZ* promoter in an *in vitro* electrophoretic mobility shift assay (EMSA) whereas LuxT-6xHis did not bind to control DNA (Fig 1D). The laddering present in the P*_swrZ_* EMSA may indicate that multiple LuxT-binding sites exist in the *swrZ* promoter and/or that LuxT oligomerizes when binding the *swrZ* promoter (Fig 1D).

### Both *luxT* and *swrZ* are subject to feedback regulation, and neither *luxT* nor *swrZ* is controlled by QS

To further explore connections between LuxT and SwrZ in *V. harveyi*, we assessed whether feedback regulatory loops exist. First, regarding SwrZ regulation of *luxT*: We find no evidence for SwrZ-mediated feedback onto *luxT* because *luxT* transcript levels were similar in WT and Δ*swrZ V. harveyi* (Fig 1E), and, moreover, overexpression of *swrZ* in WT *V. harveyi* did not alter *luxT* transcription as measured by qRT-PCR (Fig S2). qRT-PCR validated that *swrZ* was indeed overexpressed from the p*swrZ* plasmid (Fig S2). Second, we investigated autoregulation of *luxT*. Activity of a P*_luxT_-lux* reporter was measured in Δ*luxA* and Δ*luxA* Δ*luxT V. harveyi* strains. *V. harveyi* is naturally bioluminescent and *luxA* encodes a luciferase subunit. Thus, deletion of *luxA* was required to ensure that all light produced by the test strains came from the P*_luxT_*-*lux* reporter. Fig 1F shows that, in strains harboring an empty control vector, P*_luxT_-lux* expression is 6-fold higher in the Δ*luxA V. harveyi* strain than in the Δ*luxA* Δ*luxT* strain. Introduction of a vector expressing *luxT* restored light production (Fig 1F). Thus, LuxT activates its own transcription. Unfortunately, autoregulation of *luxT* could not be tested in recombinant *E. coli* because the P*_luxT_-lux* reporter was not expressed to any detectable level. However, in an *in vitro* EMSA, LuxT-6xHis bound the *luxT* promoter, providing evidence for a direct auto-regulatory role (Fig 1G). Finally, we investigated direct autoregulation of *swrZ*. In recombinant *E. coli,* induction of *swrZ* expression repressed a P*_swrZ_-lux* reporter 30-fold indicating direct feedback repression (Fig 1H). We conclude that LuxT and SwrZ can be placed into a regulatory pathway in which LuxT directly represses *swrZ.* Positive and negative feedback loops exist for LuxT and SwrZ, respectively (Fig 1I). Below, we speculate on the implications of this regulatory arrangement.

Analysis of the LuxT regulon revealed that a subset of LuxT-controlled genes are also regulated by QS ((8) and Data Set S1). One possible explanation for this finding is that regulatory interconnections exist between LuxT and QS. We know that LuxT does not regulate *aphA* or *luxR,* so LuxT cannot reside upstream of those two components in the QS cascade (18). Alternatively, QS could control *luxT* and/or *swrZ* expression. To test this possibility, we measured *luxT* and *swrZ* expression in WT, Δ*luxO,* and *luxO* D61E *V. harveyi* strains. The logic is as follows: The WT strain undergoes the normal LCD to HCD QS transition. The Δ*luxO* strain exhibits “HCD-locked” behavior because in the absence of LuxO activity, no Qrr sRNAs are produced ((9, 10) and Fig S1B). *luxO* D61E encodes a LuxO-P mimetic, so the strain harboring this mutation exhibits “LCD-locked” behavior in which the Qrr sRNAs are constitutively produced ((10, 17) and Fig S1A). There were no significant differences in *luxT* or *swrZ* expression in these strains (Fig S3A). Consistent with this finding, no dramatic changes in *luxT* expression occurred over growth as would be expected for a QS-regulated gene (Fig S3B). As expected from our above findings (Fig 1B), *swrZ* transcript levels were > 100-fold higher in the Δ*luxT* strain than in WT *V. harveyi*, but again, cell-density-dependent changes in *swrZ* expression did not occur in either strain, arguing against QS control (Fig S3C). We conclude that expression of *luxT* and *swrZ* are not QS regulated. Thus, we infer that QS and LuxT/SwrZ function independently and in parallel in the regulation of a subset of genes in each of their regulons.

### LuxT activates expression of type III secretion system genes via repression of *swrZ*

As mentioned above, LuxT regulates ∼413 genes in addition to *swrZ* (Fig 1A and Data Set S1). To gain insight into the LuxT regulon, as examples, we characterize the mechanisms by which LuxT controls genes specifying three *V. harveyi* functions: type III secretion, siderophore production, and aerolysin production. We begin with type III secretion genes. Type III secretion systems (T3SSs) are syringe-like molecular machines that ferry toxic effector proteins across bacterial inner and outer membranes and into target eukaryotic cells (25–27). In *V. harveyi* and other vibrios, type III secretion is crucial for virulence (28–30). The structural proteins comprising the *V. harveyi* T3SS are encoded in four operons (T3SS.1, T3SS.2, T3SS.3, and T3SS.4; Fig 2A) (5, 8, 31). Type III secretion is QS-regulated in *V. harveyi* (5). Specifically, T3SS genes are repressed by both AphA and LuxR. The consequence is that T3SS gene expression peaks at low- to mid-cell density when AphA levels have decreased and LuxR levels have not risen (8, 12). QS regulation of type III secretion occurs indirectly through ExsA, the master transcriptional activator of T3SS genes (Fig 2A). Both AphA and LuxR directly repress *exsA* (8, 31).

**Fig 2.**
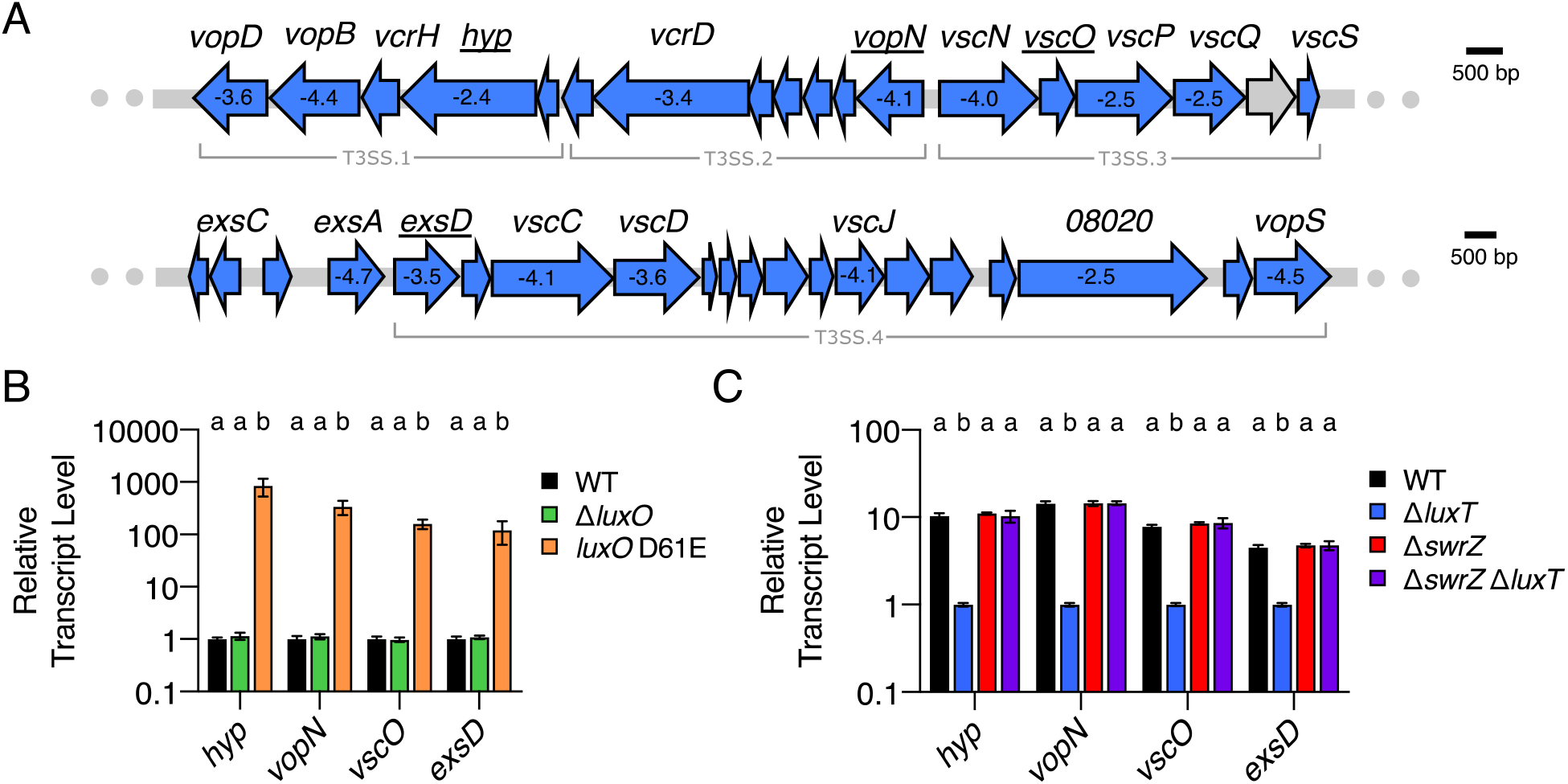
QS and LuxT/SwrZ regulate *V. harveyi* T3SS gene expression. **(A)** Schematic of T3SS gene organization in *V. harveyi.* There are four major T3SS operons, as labeled. All genes shown in blue were identified by RNA-Seq as members of the LuxT regulon. The numbers shown within select genes designate the fold-change differences in transcript levels between WT and Δ*luxT V. harveyi*, as measured by RNA-Seq (Data Set S1). Expression of underlined genes was measured in panels B and C. **(B)** qRT-PCR measurements of mRNA levels of the indicated genes in WT (black), Δ*luxO* (green), and *luxO* D61E (orange) *V. harveyi* strains. RNA was isolated from strains grown in AB to OD_600_ = 1. **(C)** qRT-PCR measurements of mRNA levels of the indicated genes in WT (black), Δ*luxT* (blue), Δ*swrZ* (red), and Δ*swrZ* Δ*luxT* (purple) *V. harveyi* strains. RNA was isolated from strains grown in LM to OD_600_ = 0.1. In B and C, error bars represent standard deviations of the means of *n* = 3 biological replicates. Different letters indicate significant differences between strains, *p* < 0.05 (two-way analysis of variation (ANOVA) followed by Tukey’s multiple comparisons test).

The genes in all four T3SS operons and *exsA* were revealed by RNA-Seq as activated by LuxT (Data Set S1 and Fig 2A). To understand dual control of T3SS genes by QS and LuxT, we engineered a set of strains and developed a companion strategy to probe regulatory mechanisms. First, we verified LuxT involvement from the RNA-Seq analysis, and we determined if *qrr*1 is required for LuxT control of T3SS genes. Using qRT-PCR, we quantified transcript levels of four T3SS genes: *hyp, vopN, vscO,* and *exsD*, representing the four T3SS operons (underlined in Fig 2A). Expression was compared between WT and Δ*luxT V. harveyi* and between Δ*qrr*1 and Δ*qrr*1 Δ*luxT V. harveyi*. Expression was, respectively, ∼10, 14, 8, and 4-fold lower in the Δ*luxT* strain than in WT, confirming that LuxT is an activator of T3SS genes (Fig S4A). Nearly identical results to those shown in Fig S4A were obtained for the Δ*qrr*1 and Δ*qrr*1 Δ*luxT* strains, indicating that LuxT activates T3SS genes independently of Qrr1 (Fig S4B).

To verify the mechanism by which QS regulates T3SS genes, we measured transcript levels of the four representative genes in WT, Δ*luxO* (HCD-locked), and *luxO* D61E (LCD-locked) *V. harveyi* strains. Expression of *hyp, vopN, vscO,* and *exsD* were > 100-fold higher in the *luxO* D61E strain than in the WT and Δ*luxO* strains at HCD (Fig 2B). Consistent with previously published results, high level expression of T3SS genes occurs at LCD due to LuxR-mediated repression of them at HCD (5, 8). Specifically, compared to WT, a Δ*luxR V. harveyi* strain exhibited > 100-fold higher expression of the four genes (Fig S5A). AphA is known to repress T3SS.1 and T3SS.4 in *V. harveyi* (8). Expression of *hyp* (T3SS.1) and *exsD* (TS33.4) were ∼1.6-fold higher in the *luxO* D61E Δ*aphA* strain compared to the *luxO* D61E strain, however, the differences were not statistically significant (Fig S5B). We presume that we did not observe a larger role for AphA here because of differences in our experimental growth conditions compared to those used previously and the use of RNA-seq compared to microarrays (8).

To assess the mechanism by which LuxT controls T3SS genes expression, we first determined whether LuxT-mediated activation of T3SS genes requires SwrZ by measuring transcript levels of the four representative T3SS genes in WT, Δ*luxT,* Δ*swrZ,* and Δ*swrZ* Δ*luxT V. harveyi* strains at LCD. Comparison of expression levels between the WT and Δ*luxT* strains validates LuxT activation of the genes. Comparison of expression levels between the Δ*swrZ* and Δ*swrZ* Δ*luxT* strains tests whether LuxT activation of genes requires SwrZ. As above (Fig S4A), expression of *hyp, vopN, vscO,* and *exsD* was lower in the Δ*luxT* strain than in WT (Fig 2C). However, elimination of *luxT* did not alter transcript levels in the Δ*swrZ* background (Fig 2C). Thus, LuxT activates expression of all four T3SS operons via a SwrZ-dependent mechanism. Most likely, LuxT represses *swrZ,* and SwrZ represses T3SS genes. We test this assumption below.

### LuxT regulates *exsA* by SwrZ-dependent and SwrZ-independent mechanisms to activate type III secretion

To quantify the contributions of QS and LuxT/SwrZ to regulation of type III secretion, we used Western blot assessment of VopD, a T3SS.1 protein that accumulates in and is secreted from the cytoplasm following activation of T3SS gene expression (5). We measured VopD levels in WT, Δ*luxT,* Δ*swrZ,* and Δ*swrZ* Δ*luxT V. harveyi* strains at HCD and in the LCD-locked (*luxO* D61E) strain. No VopD was detected in the strains at HCD, consistent with LuxR repression of T3SS genes (Fig 3A, first four lanes) (5, 31). By contrast, VopD was made in the *luxO* D61E strain, and deletion of *luxT* in the *luxO* D61E strain reduced VopD levels to below our detection limit (Fig 3A, fifth and sixth lanes). These data mirror our qRT-PCR results from Fig 2C and confirm the role of LuxT as an activator of type III secretion. To assess whether *swrZ* is required for LuxT activation of type III secretion, VopD levels were measured in *luxO* D61E Δ*swrZ* and *luxO* D61E Δ*swrZ* Δ*luxT* strains. Lower levels of VopD were present in the *luxO* D61E Δ*swrZ* Δ*luxT* strain than in the *luxO* D61E Δ*swrZ* strain (Fig 3A, right-most two lanes). These results differ from the qRT-PCR results in Fig 2C, likely because of the different growth conditions required for the qRT-PCR and the VopD Western blot analyses. We summarize the data in Fig 3A as follows: Deletion of *luxT* reduces VopD production in the presence and absence of *swrZ,* however a more dramatic reduction occurs when *swrZ* is present. To explain these data, we developed and tested the following model for LuxT/SwrZ regulation of T3SS genes:

**Fig 3.**
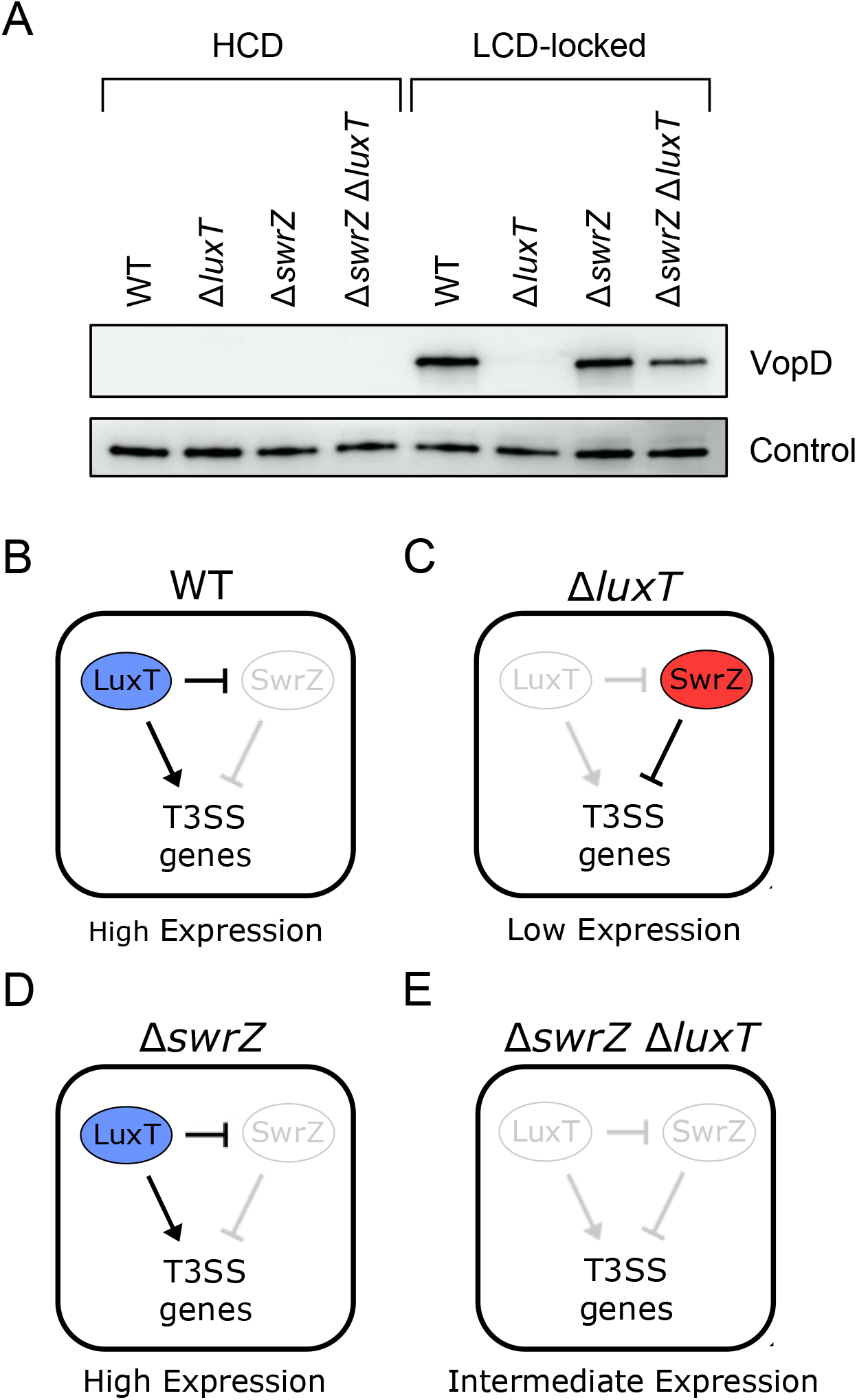
LuxT activates *V. harveyi* type III secretion gene expression by two mechanisms. **(A)** Western blot of cytoplasmic VopD levels in the indicated *V. harveyi* strains at HCD. “LCD-locked” denotes that the parent strain for the samples in the right four lanes harbors the *luxO* D61E mutation. Detection of LuxS protein serves as the loading control, as previously described (5). **(B-E)** Schematics depicting the outcomes for LuxT regulation of T3SS genes by SwrZ-dependent and SwrZ-independent mechanisms for WT **(B),** Δ*luxT* **(C),** Δ*swrZ* **(D)**, and Δ*swrZ* Δ*luxT* **(E)** scenarios. See text for details.

We propose that LuxT activates T3SS genes by two mechanisms: One mechanism is SwrZ-dependent, and one mechanism is SwrZ-independent. This model is depicted in Fig 3B-E and is used to explain the data in the LCD-locked strains of the Western blot shown in Fig 3A (right-most four lanes). In WT *V. harveyi* (Fig 3B), LuxT is present. It represses *swrZ,* and it also activates T3SS genes by a SwrZ-independent mechanism. The consequence is LuxT activation of T3SS genes. Therefore, high-level VopD production occurs (fifth lane in Fig 3A). In Δ*luxT V. harveyi* (Fig 3C), *swrZ* is derepressed. The consequence is that SwrZ is produced and available to repress T3SS genes. Therefore, no VopD is produced (sixth lane in Fig 3A). In the Δ*swrZ* strain (Fig 3D), the situation is essentially identical to WT in which *swrZ* is repressed by LuxT, but in this case, there simply is no *swrZ.* The consequence is LuxT activation of T3SS genes. Therefore, high-level VopD production occurs (seventh lane in Fig 3A). Finally, in the case of the Δ*swrZ* Δ*luxT* double mutant (Fig 3E), there is no repression of T3SS genes by SwrZ and there is no activation by LuxT. Thus, basal level expression of T3SS genes ensues, and an intermediate amount of VopD is made (eighth lane in Fig 3A). Key to this model is that VopD levels in the LCD-locked Δ*swrZ* and Δ*swrZ* Δ*luxT* strains are not identical because of the SwrZ-independent mechanism by which LuxT activates T3SS genes (Fig 3D and 3E).

To test our model, we assessed whether LuxT activates T3SS gene expression by a SwrZ-independent mechanism. To do this, we introduced a p*luxT* overexpression vector into a Δ*luxR* Δ*swrZ* Δ*luxT V. harveyi* strain. The Δ*luxR* mutation was included to eliminate HCD repression of T3SS genes. This strategy is superior to making measurements from RNA collected at LCD from strains harboring *luxR*. In the latter case, residual LuxR repressor is present (see Fig S5A). In this setup, we measured *hyp, vopN, vscO,* and *exsD* transcript levels. We also measured expression of *exsA,* which is located immediately upstream of T3SS.4 and, as mentioned, encodes the master transcriptional activator of T3SS genes. Compared to the vector control, overexpression of *luxT* caused ∼122, 74, 34, 57, and 19-fold increases in *hyp, vopN, vscO, exsD,* and *exsA* expression, respectively (Fig 4A). These data demonstrate that LuxT activates the four T3SS operons and *exsA* by a mechanism that does not require SwrZ. We also assessed *swrZ* overexpression in this experimental setup. Repression by SwrZ was not observed (Fig 4A) presumably because in the absence of LuxT activation, T3SS gene expression is minimal so no detectable repression by SwrZ can occur. To test this supposition, we introduced the p*swrZ* overexpression construct into a Δ*luxR* Δ*swrZ V. harveyi* strain. Indeed, in the *luxT^+^* background, overexpression of *swrZ* caused an ∼10-fold reduction in *hyp, vopN,* and *vscO* transcript levels and ∼9 and 3-fold reductions in *exsD* and *exsA* levels, respectively (Fig 4B). Thus, SwrZ is a repressor of the four T3SS operons and *exsA*.

**Fig 4.**
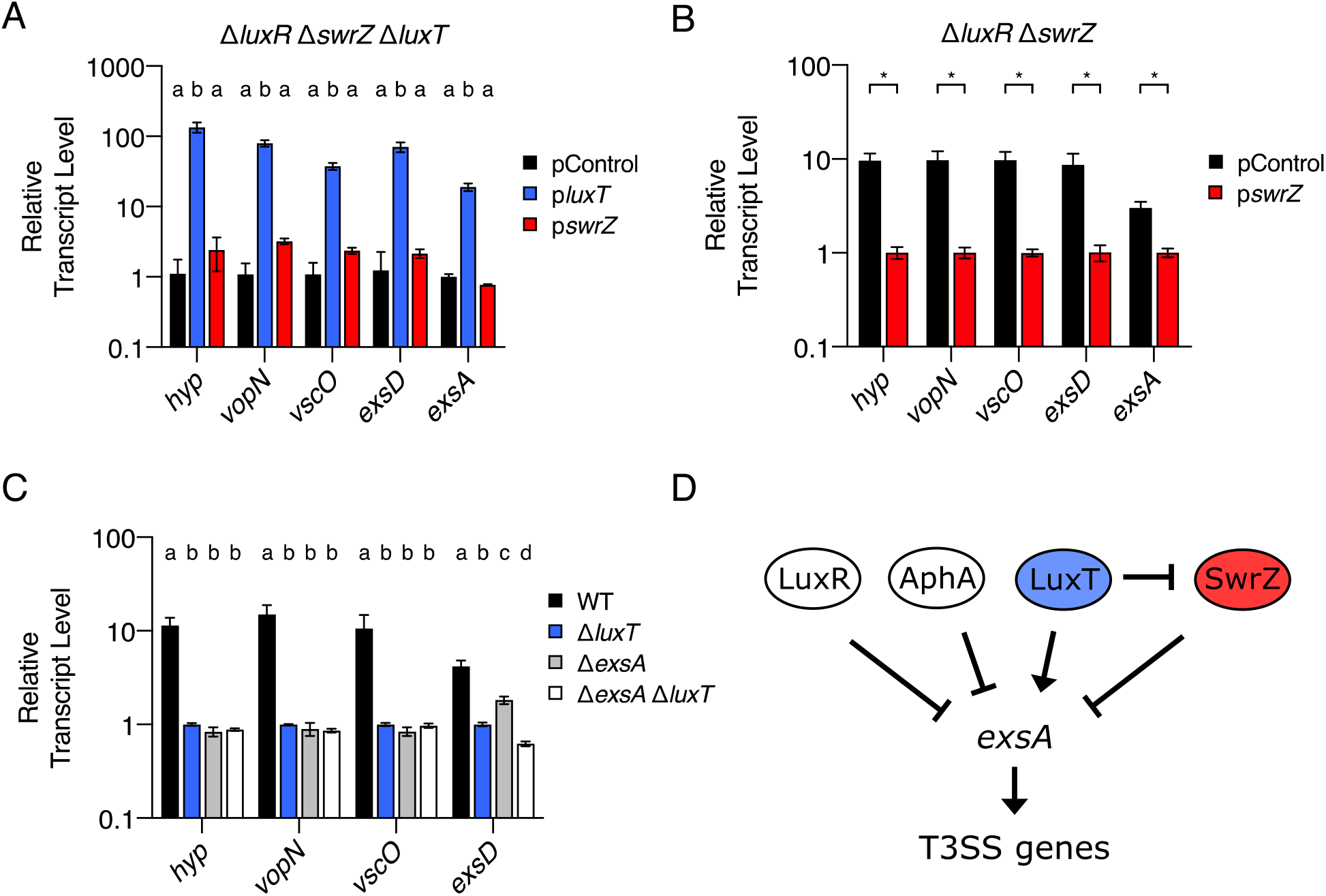
LuxT activates *V. harveyi exsA*, and in turn, T3SS genes, by SwrZ-dependent and SwrZ-independent mechanisms. **(A)** qRT-PCR measurements of transcript levels of the indicated genes in Δ*luxR* Δ*swrZ* Δ*luxT V. harveyi* harboring the indicated plasmid. pControl (black) denotes the empty parent vector, and p*luxT* (blue) and p*swrZ* (red) denote vectors harboring IPTG-inducible *luxT* and *swrZ,* respectively. RNA was isolated from strains grown for 16 h in AB supplemented with 0.5 mM IPTG. **(B)** As in A, for Δ*luxR* Δ*swrZ V. harveyi* harboring the indicated plasmid. **(C)** qRT-PCR measurements of transcript levels of the indicated genes in WT (black), Δ*luxT* (blue), Δ*exsA* (gray), and Δ*exsA* Δ*luxT* (white) *V. harveyi* strains. RNA was isolated from *V. harveyi* strains grown in LM to OD_600_ = 0.1. **(D)** Model for QS and LuxT/SwrZ regulation of T3SS genes. In A- C, error bars represent standard deviations of the means of *n* = 3 biological replicates. In A and C, different letters indicate significant differences between strains, *p* < 0.05 (two- way ANOVA followed by Tukey’s multiple comparisons test). In B, unpaired two-tailed *t* tests with Welch’s correction were performed comparing the indicated pControl and p*swrZ* conditions. *p*-value: * < 0.05.

Previously, it was discovered that T3SS gene expression does not occur in the absence of *exsA* irrespective of the presence or absence of LuxR. Thus, ExsA is epistatic to QS in the control of T3SS genes (31). Consistent with this finding, overexpression of *exsA* overrode repression by LuxR to activate T3SS gene expression at HCD (31). Based on these earlier data and our identification of LuxT and SwrZ as regulators of T3SS genes, we wondered whether LuxT and SwrZ also exert control over T3SS genes via regulation of *exsA*. To test this possibility, we measured *hyp, vopN, vscO,* and *exsD* expression in WT, Δ*luxT*, Δ*exsA,* and Δ*exsA* Δ*luxT V. harveyi* strains. Fig 4C shows that, as expected, expression of all four genes was lower in the Δ*luxT* strain than in WT. However, there were no significant differences in *hyp, vopN,* and *vscO* transcript levels between the Δ*exsA* and Δ*exsA* Δ*luxT* strains (Fig 4C). We conclude that *exsA* is required for LuxT activation of T3SS.1, T3SS.2, and T3SS.3, encompassing *hyp, vopN,* and *vscO,* respectively. Because LuxT activation of *hyp, vopN,* and *vscO* requires *swrZ* (see Fig 2C), we can also conclude that regulation by SwrZ requires *exsA*. Regarding T3SS.4, there was ∼3-fold less *exsD* transcript present in the Δ*exsA* Δ*luxT* strain than the Δ*exsA* strain. It is possible that LuxT activates the T3SS.4 operon encompassing *exsD* independently of ExsA or, alternatively, based on their sequential orientation in the genome (Fig 2A), LuxT only activates the *exsA* promoter and read-through transcription occurs for T3SS.4.

Together, our results show that LuxT activates T3SS genes by two mechanisms; one is SwrZ-dependent, and one is SwrZ-independent. Also, LuxT functions independently of QS to control these genes. QS and LuxT/SwrZ all modulate T3SS.1, T3SS.2, and T3SS.3 via control of expression of *exsA* encoding a transcriptional activator of T3SS genes. Previous results showed that *exsA* is repressed by both AphA and LuxR, which constrains its expression to low- to mid-cell densities. Our scheme for QS and LuxT/SwrZ regulation of T3SS genes is depicted in Fig 4D.

### QS and LuxT/SwrZ regulate *V. harveyi* siderophore production

Similar to type III secretion, the RNA-Seq analysis revealed LuxT to be an activator of siderophore production genes in *V. harveyi.* Siderophores are small molecule iron chelators that bacteria produce and secrete to scavenge extracellular iron (32). Companion siderophore uptake systems import siderophore-Fe^3+^ complexes, facilitating iron acquisition. *V. harveyi* encodes the genes required to produce, secrete, and import two siderophores: amphi-enterobactin and anguibactin (33, 34), and both sets of genes are regulated by LuxT (Fig 5A and 5B, respectively, and Data Set S1). *V. harveyi* produces siderophores at LCD suggesting QS regulation, however, the mechanism connecting QS to siderophore genes is not defined (6, 35). Using the steps laid out above for characterization of T3SS gene regulation, we determine how QS, LuxT, and SwrZ control siderophore production.

**Fig 5.**
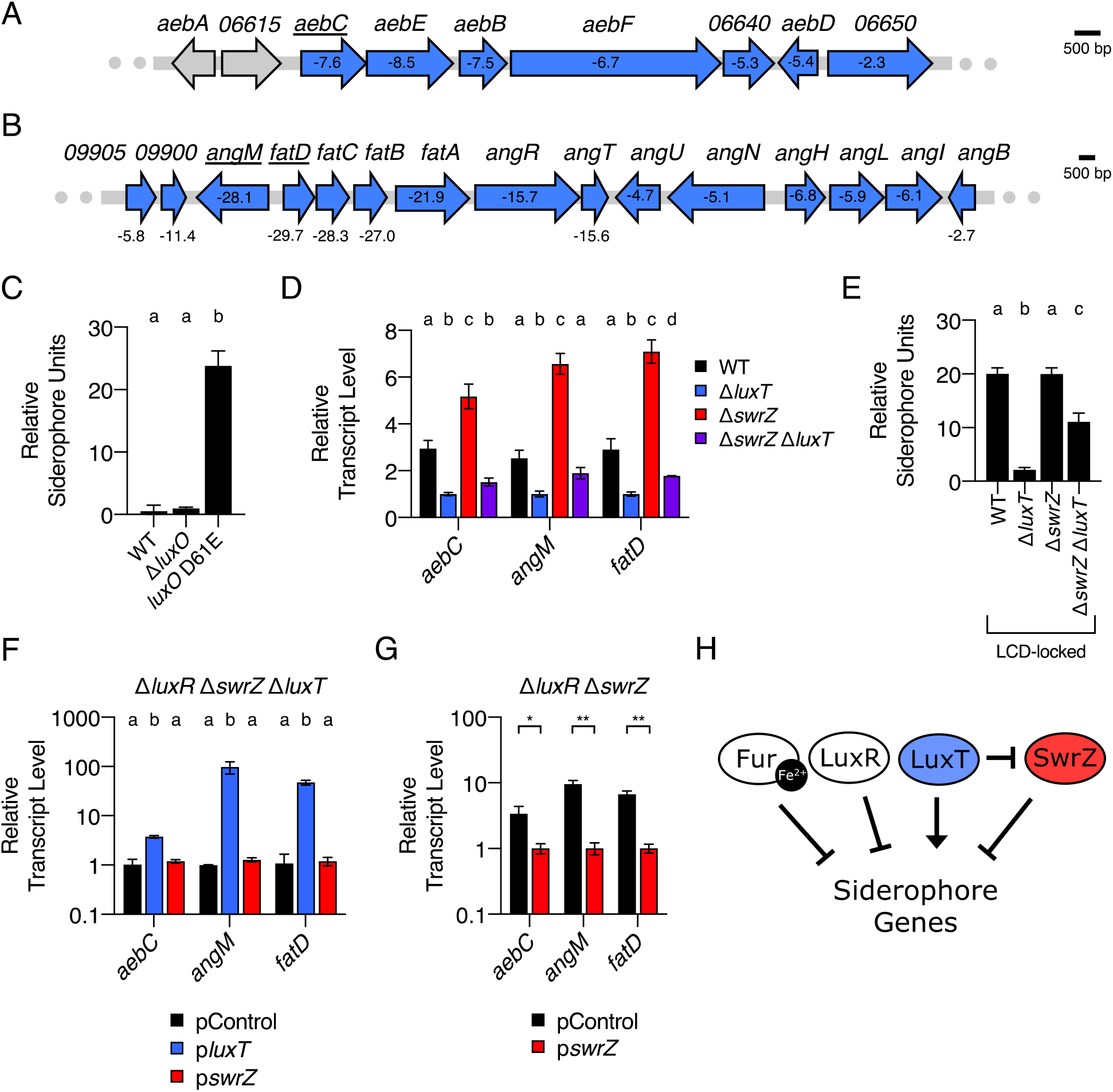
QS, LuxT, and SwrZ regulate *V. harveyi* siderophore production. **(A-B)** Schematic of siderophore gene organization in *V. harveyi* for amphi-enterobactin (A) and anguibactin (B). All genes labeled in blue were identified by RNA-Seq as members of the LuxT regulon. The numbers shown within or below the genes designate the fold-change differences in transcript levels between WT and Δ*luxT V. harveyi*, as measured by RNA- Seq. Expression of underlined genes was measured in panels D, F, and G. **(C)** CAS assay quantitation of siderophore levels in cell-free culture fluids isolated from the indicated *V. harveyi* strains. Strains were grown for 16 h in AB medium. **(D)** qRT-PCR measurements of transcript levels of the indicated genes in WT (black), Δ*luxT* (blue), Δ*swrZ* (red), and Δ*swrZ* Δ*luxT* (purple) *V. harveyi* strains. RNA was isolated from strains grown in LM to OD_600_ = 0.1. **(E)** As in C. “LCD-locked” denotes that the strains harbor the *luxO* D61E mutation. **(F)** qRT-PCR measurements of transcript levels of the indicated genes in Δ*luxR* Δ*swrZ* Δ*luxT V. harveyi* harboring the indicated plasmid. pControl (black) denotes the empty parent vector, and p*luxT* (blue) and p*swrZ* (red) denote vectors harboring IPTG-inducible *luxT* and *swrZ,* respectively. RNA was isolated from strains grown for 16 h in AB supplemented with 0.5 mM IPTG. **(G)** As in F, for Δ*luxR* Δ*swrZ V. harveyi* harboring the indicated plasmid. **(H)** Model for Fur, QS, and LuxT/SwrZ regulation of *V. harveyi* siderophore production. In C-G, error bars represent standard deviations of the means of *n* = 3 biological replicates. In C-F, different letters indicate significant differences between strains, *p* < 0.05 (two-way ANOVA followed by Tukey’s multiple comparisons test). In G, unpaired two-tailed *t* tests with Welch’s correction were performed comparing the indicated pControl and p*swrZ* conditions. *p*-values: * < 0.05, ** < 0.01.

First, qRT-PCR validated the RNA-Seq data. Transcript levels of *aebC* encoding isochorismate synthase were measured as a representative amphi-enterobactin biosynthetic gene (33). Regarding anguibactin, expression of *angM* and *fatD* were measured, and they encode a non-ribosomal peptide synthetase and an anguibactin-Fe^3+^ transporter, respectively (36, 37). Transcript levels of *aebC, angM,* and *fatD* were ∼3-fold lower in Δ*luxT V. harveyi* than in WT (Fig S4C). Analogous results were obtained for the three genes when measured in Δ*qrr*1 and Δ*qrr*1 Δ*luxT* strains, showing that LuxT activates siderophore genes independently of Qrr1 (Fig S4D).

Second, to determine the mechanism by which QS controls siderophore production, we used a chrome azurol S (CAS) dye to quantify siderophore levels in cell-free culture fluids prepared from WT *V. harveyi* and QS mutant strains. Approximately 24- fold more siderophore was present in fluids from the LCD-locked *luxO* D61E strain than in fluids from the WT and Δ*luxO* strains, consistent with the known LCD production pattern (Fig 5C) (6, 35). We used mutants defective in each siderophore biosynthesis pathway to show that the siderophore detected by the CAS assay was amphi-enterobactin, not anguibactin as follows: AebF is required for amphi-enterobactin production (33), and the *luxO* D61E Δ*aebF* strain had almost no siderophore in its cell-free culture fluids (Fig S6A). In contrast, the *luxO* D61E Δ*angN* strain, which lacks a gene required for anguibactin production (38), did not exhibit reduced siderophore production compared to the *luxO* D61E strain (Fig S6A). *V. harveyi* apparently does not produce anguibactin when grown under standard laboratory conditions as reported previously (35).

With respect to *V. harveyi* QS, three mechanisms can explain how a gene acquires a LCD expression pattern (Fig S1): It is activated at LCD by AphA, it is activated, directly or indirectly, at LCD by the Qrr sRNAs, or it is repressed at HCD by LuxR. Regarding AphA: Siderophore levels were similarly high in fluids from the *luxO* D61E and *luxO* D61E Δ*aphA V. harveyi* strains, so AphA has no role in regulation of siderophore production (Fig S6B). Regarding LuxR: The Δ*luxR* strain possessed ∼16-fold more amphi-enterobactin in its culture fluids than did WT at HCD (Fig S6C). Thus, QS control of amphi-enterobactin production occurs via LuxR repression at HCD. Regarding the Qrr sRNAs: At HCD somewhat less amphi-enterobactin was produced by the Δ*luxR* strain than by the *luxO* D61E strain (Fig S6B and S6C), hinting that, in addition to the primary repressive role of LuxR at HCD, at LCD, the Qrr sRNAs could activate siderophore biosynthesis or export. Possibilities include direct post-transcriptional activation by the Qrr sRNAs or alternatively, another QS-regulated factor may exist that is involved in amphi-enterobactin gene regulation. We did not investigate this mechanism further.

To position LuxT and SwrZ in the siderophore regulatory pathway, we used qRT-PCR quantitation of *aebC, angM,* and *fatD* in WT, Δ*luxT,* Δ*swrZ,* and Δ*swrZ* Δ*luxT V. harveyi* strains at LCD (Fig 5D). As in Fig S4C, there were lower mRNA levels in the Δ*luxT* strain than in the WT showing again that LuxT activates siderophore genes. Modestly increased expression of the three genes occurred in the Δ*swrZ* strain compared to the WT, indicating that SwrZ is a repressor of siderophore genes. Finally, the Δ*luxT* Δ*swrZ* double mutant expressed siderophore genes at a level intermediate between those of the Δ*luxT* and Δ*swrZ* strains. These data resemble the findings in Fig 3A for LuxT/SwrZ regulation of T3SS genes, suggesting an identical model: LuxT activates siderophore production by both a SwrZ-dependent and a SwrZ-independent mechanism. We validate this model below.

### Fur represses *V. harveyi* siderophore production in iron-rich conditions

The above qRT-PCR measurements of siderophore transcripts showed only modest differences between strains (Fig 5D) possibly as a consequence of the generally low siderophore gene expression that occurs in iron-replete conditions due to repression by Fur, the major transcriptional regulator of iron transport. Fur repression is relieved in iron-limiting conditions (39). Indeed, culture fluids used for the CAS assays in Fig 5C and S6A-C were prepared from *V. harveyi* grown in low-iron minimal medium, and significant siderophore was produced. To assess the role of Fur in regulation of *V. harveyi* siderophore production, *aebC, angM,* and *fatD* transcript levels were compared between WT and Δ*fur V. harveyi*. In iron-rich medium, in the absence of *fur*, expression of the three test genes was ∼284, 119, and 66-fold higher than when *fur* was present (Fig S6D). Deletion of *luxT* in the Δ*fur* strain caused 3, 43, and 52-fold decreases in *aebC, angM,* and *fatD* transcript levels, respectively (Fig S6D). Thus, Fur is a repressor of amphi- enterobactin and anguibactin genes, which diminished our ability to observe their regulation by LuxT/SwrZ in iron-replete conditions, particularly for *angM* and *fatD.* We note that *fur* expression is not controlled by LuxT or SwrZ (Fig S6E). Thus, Fur and LuxT/SwrZ regulate siderophore genes independently. The siderophore experiments in the next section were conducted following growth of *V. harveyi* under iron-limited conditions.

### LuxT activates *V. harveyi* siderophore production by SwrZ-dependent and SwrZ- independent mechanisms

To validate the roles of and relationship between LuxT and SwrZ in regulation of siderophore production, we quantified siderophore produced by the LCD-locked *luxO* D61E, *luxO* D61E Δ*luxT*, *luxO* D61E Δ*swrZ*, and *luxO* D61E Δ*swrZ* Δ*luxT* mutant strains using the CAS assay. Consistent with the qRT-PCR results (Fig 5D), the *luxO* D61E Δ*swrZ* Δ*luxT* mutant produced intermediate siderophore levels between those of the *luxO* D61E Δ*luxT* strain and the *luxO* D61E Δ*swrZ* strain (Fig 5E). These data support a model in which LuxT activates siderophore production by two mechanisms – one dependent on SwrZ and one independent of SwrZ. We performed complementation analyses in a *V. harveyi* Δ*luxR* Δ*luxT* Δ*swrZ* strain as a final test of our model. Even in the absence of *swrZ,* overexpression of *luxT* caused increased expression of *aebC, angM,* and *fatD,* supporting the SwrZ-independent role for LuxT (Fig 5F). Following the reasoning described above for T3SS genes, repression due to *swrZ* overexpression could only be observed in the *luxT*^+^ background (Fig 5F and 5G). Our model for regulation of siderophore production in *V. harveyi* is depicted in Fig 5H: Fur represses siderophore genes in iron-replete conditions, and LuxR represses siderophore genes at HCD. LuxT activates siderophore genes by both a SwrZ-dependent and a SwrZ-independent mechanism.

### QS and LuxT/SwrZ regulate production of the aerolysin toxin

The final *V. harveyi* LuxT-regulated behavior we investigated from the RNA-Seq was production of the pore-forming toxin aerolysin. Aerolysin-like proteins, originally discovered in the bacterium *Aeromonas hydrophila,* are a family of secreted proteins that form transmembrane β-barrel pores in eukaryotic target cells, causing cell death (40–42). Aerolysins are virulence factors in *Aeromonas* spp. and in *Vibrio splendidus* (43, 44).

We previously reported that LuxT is an activator of *VIBHAR_RS11620* encoding an aerolysin toxin. For clarity, we now call *VIBHAR_RS11620 aerB.* We determined that LuxT activates *aerB* expression by two mechanisms. First, LuxT activates *aerB* transcription by a Qrr1-independent mechanism. Second, LuxT indirectly activates *aerB* translation by repressing expression of *qrr*1 encoding a post-transcriptional repressor of *aerB* (13, 18)*. V. harveyi* aerolysin production can be measured by monitoring hemolysis of blood cells in liquid or on blood agar plates. Activation by LuxT is required for *V. harveyi* hemolytic activity in both assays. Observable repression by Qrr1 occurs only in the plate assay (18).

In agreement with our earlier findings, the RNA-Seq revealed LuxT as an activator of the *aerB* aerolysin toxin gene. Three additional genes upstream of *aerB* were also activated (Fig 6A and Data Set S1): *VIBHAR_RS11600*, which we call *aerR*, encodes a transcriptional regulator, *VIBHAR_RS11605*, which we do not name, encodes a protein of unknown function, and *VIBHAR_RS11610*, which we call *aerA*, encodes an additional aerolysin family toxin. The protein sequences of AerA and AerB are similar (50% pairwise identity, 66% pairwise positive), indicating a possible gene duplication event. The *VIBHAR_RS11615* gene that resides between *aerA* and *aerB* was not identified as regulated by LuxT (Fig 6A).

**Fig 6.**
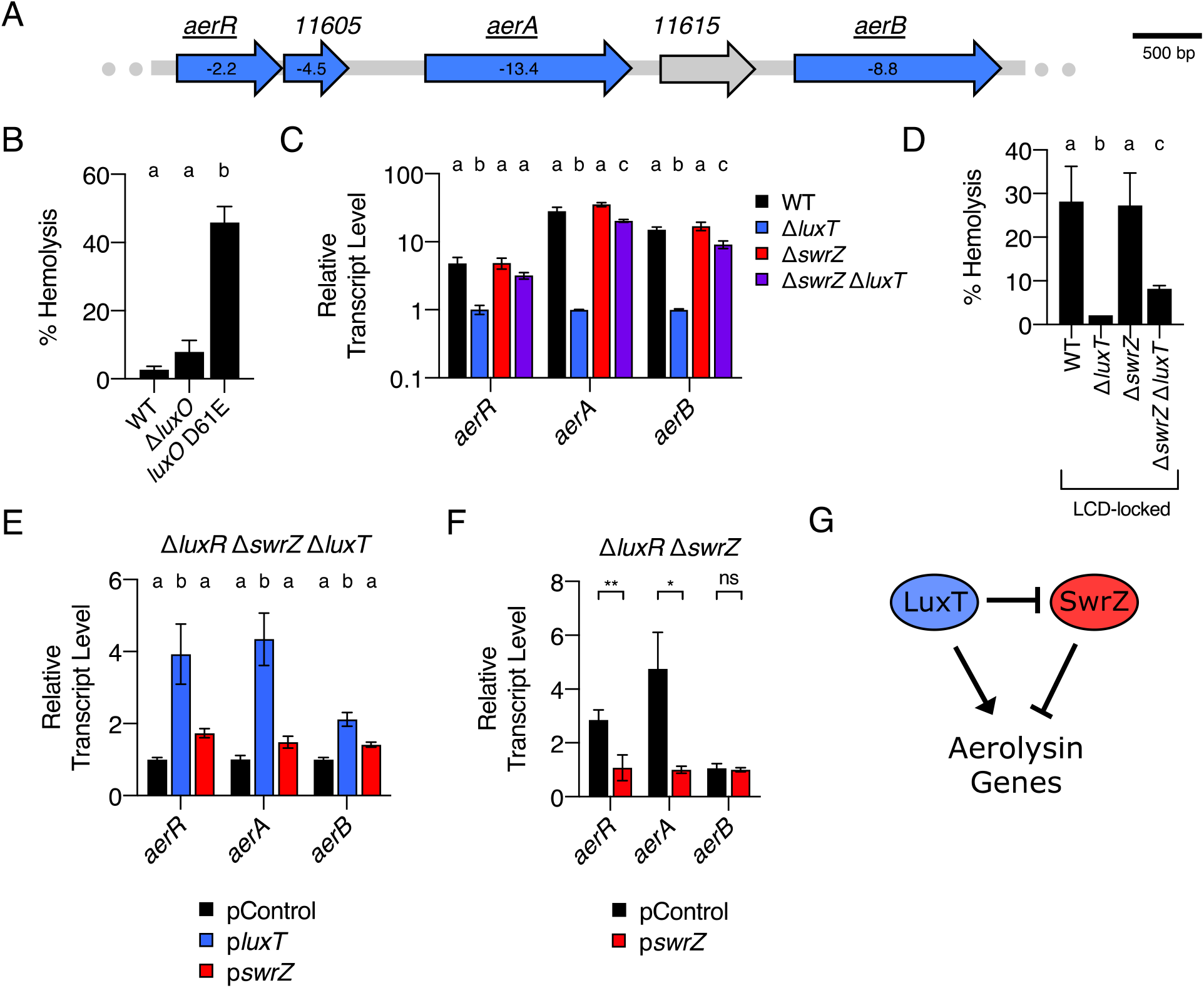
QS, LuxT, and SwrZ regulate *V. harveyi* aerolysin production. **(A)** Schematic of aerolysin gene organization in *V. harveyi*. All genes labeled in blue were identified by RNA-Seq as members of the LuxT regulon. The numbers shown within the genes designate the fold-change differences in transcript levels between WT and Δ*luxT V. harveyi*, as measured by RNA-Seq. Expression of underlined genes was measured in panels C, E, and F. **(B)** Hemolytic activity in the indicated *V*. *harveyi* cell-free culture fluids as judged by lysis of defibrinated sheep’s blood cells. Culture fluids were collected after 24 h of growth in AB medium. **(C)** qRT-PCR measurements of transcript levels of the indicated genes in WT (black), Δ*luxT* (blue), Δ*swrZ* (red), and Δ*swrZ* Δ*luxT* (purple) *V. harveyi* strains. RNA was isolated from strains grown in LM to OD_600_ = 0.1. **(D)** As in B for the designated strains. **(E)** qRT-PCR measurements of transcript levels of the indicated genes in Δ*luxR* Δ*swrZ* Δ*luxT V. harveyi* harboring the indicated plasmid. pControl (black) denotes the empty parent vector, and p*luxT* (blue) and p*swrZ* (red) denote vectors harboring IPTG-inducible *luxT* and *swrZ,* respectively. RNA was isolated from strains grown for 16 h in AB supplemented with 0.5 mM IPTG. **(F)** As in E, for Δ*luxR* Δ*swrZ V. harveyi* harboring the indicated plasmid. **(G)** Model for LuxT/SwrZ regulation of aerolysin production. In B-F, error bars represent standard deviations of the means of *n* = 3 biological replicates. In B-E, different letters indicate significant differences between strains, *p* < 0.05 (two-way ANOVA followed by Tukey’s multiple comparisons test). In F, unpaired two-tailed *t* tests with Welch’s correction were performed comparing the indicated pControl and p*swrZ* conditions. *p*-values: ns ≥ 0.05, * < 0.05, ** < 0.01.

As above, we first confirmed LuxT regulation of *aerR, aerA,* and *aerB*, and determined if Qrr1 is required. Deletion of *luxT* in WT *V. harveyi* resulted in ∼5, 28, and 15-fold lower expression of *aerR, aerA,* and *aerB,* respectively (Fig S4E). In a strain lacking *qrr*1, deletion of *luxT* caused similar reductions in expression (Fig S4F). These data show that Qrr1 is not required for LuxT activation of aerolysin genes. We know from our earlier work that Qrr1 is a post-transcriptional regulator of *aerB.* Regulation by Qrr1 can only be detected when measuring *aerB* translation, not transcription (13, 18). For the remainder of the present work, we focus on the transcriptional, Qrr1-independent, mechanism by which LuxT activates aerolysin production.

QS regulation of aerolysin production is more complicated than we anticipated, nonetheless, we lay out what we know here. In the liquid hemolysis assay, culture fluids from the LCD-locked *luxO* D61E strain possessed ∼17-fold more hemolysis activity than fluids from HCD WT *V. harveyi* (Fig 6B). Fluids from the HCD-locked Δ*luxO* strain had activity similar to that of the WT strain (Fig 6B). Thus, aerolysin production is QS-controlled and produced at LCD. Deletion of *aphA* in the *luxO* D61E LCD-locked strain did not reduce hemolysis (Fig S7A), and deletion of *luxR* in the WT did not increase hemolysis at HCD (Fig S7B). These data were unexpected and show that, while aerolysin is QS-controlled, neither AphA nor LuxR regulates its production. We confirmed the results using blood agar plates. The results on the plates were identical to those in liquid with the exception that the Δ*luxR* strain caused less hemolysis on plates than did the WT (Fig S7C). Given the LCD production pattern for aerolysin, we conclude that aerolysin genes must be activated at LCD by the QS Qrr sRNAs, either directly or indirectly. We note that this finding seemingly contradicts our earlier result showing that Qrr sRNAs post-transcriptionally repress *aerB* (13, 18). We preliminarily predict that the Qrr sRNAs exert both positive and negative effects on aerolysin production. Activation by the Qrr sRNAs is likely indirect through an unknown regulator, while post-transcriptional repression of *aerB* occurs directly (13). Further investigation to characterize the mechanism of QS control of aerolysin genes is required.

While the mechanism underlying QS regulation of aerolysin genes is not fully defined, what is clear is that *aerR*, *aerA*, and *aerB* are all regulated by LuxT (Data Set S1 and Fig S4E). Thus, we next explored which of the three genes is required for *V. harveyi* hemolysis activity. We deleted each gene individually in the LCD-locked *luxO* D61E *V. harveyi* strain. Deletion of *aerR* and *aerA* eliminated hemolysis activity in the liquid and plate assays (Fig S7D and S7E, respectively). In contrast, deletion of *aerB* did not reduce hemolytic activity, so *aerB* is not required (Fig S7D and S7E). We speculate that AerR is an activator of *aerA,* and AerA is the primary secreted aerolysin toxin. Because *aerB* is repressed by the Qrr sRNAs, *aerB* expression may remain low at LCD.

To understand the mechanism underlying LuxT activation of aerolysin gene expression, we measured *aerR, aerA,* and *aerB* transcript levels in WT, Δ*luxT,* Δ*swrZ,* and Δ*swrZ* Δ*luxT V. harveyi* strains at LCD. Expression of all three genes was high in the WT and Δ*swrZ* strains, low in the Δ*luxT* strain, and intermediate in the Δ*swrZ* Δ*luxT* strain (Fig 6C). The corresponding hemolysis patterns in the LCD-locked *luxO* D61E background exactly mirrored the transcription patterns both in the liquid assay (Fig 6D) and on the blood agar plates (Fig S7F). Exactly analogous to what we have explained above, these data indicate that LuxT activates aerolysin production by both a SwrZ- dependent and a SwrZ-independent mechanism. We verified this assumption using complementation analyses (Fig 6E and 6F). The results were as expected according to our putative mechanism except that *swrZ* overexpression did not repress *aerB* (Fig 6F). However, the qRT-PCR results in Fig 6C show that SwrZ is a repressor of *aerB.* We presume *swrZ* complementation did not repress *aerB* because *aerB* exhibits quite low basal expression. Our model for LuxT/SwrZ regulation of aerolysin production is depicted in Fig 6G.

### *luxT* is and *swrZ* is not conserved among *Vibrionaceae* family members

As mentioned, SwrT (LuxT) repression of *swrZ* was originally discovered in *V. parahaemolyticus* and shown to be relevant for regulation of swarming motility (22). Here, we have shown that LuxT also regulates *swrZ* in *V. harveyi,* and LuxT and SwrZ both regulate type III secretion, siderophore production, and aerolysin production. LuxT also regulates genes in *V. harveyi* that SwrZ does not control. For example, LuxT activates *luxCDABE*, encoding luciferase, and SwrZ plays no role ((18, 20) and Fig S8). We assessed SwrZ involvement in the regulation of 5 additional LuxT-activated and 3 additional LuxT-repressed genes. SwrZ regulated all 5 of the LuxT-activated test genes, whereas SwrZ did not regulate the 3 LuxT-repressed test genes (Fig S8). Based on the opposing roles of LuxT and SwrZ (See Fig 3B-E), we speculate that SwrZ may only function as a transcriptional repressor, possibly explaining why it is not required to participate in LuxT repression of gene expression. We conclude that within the LuxT regulon, a subset of genes does not employ SwrZ in regulation.

In *V. harveyi* and *V. parahaemolyticus,* the *luxT* (*swrT*) and *swrZ* genes are encoded on the two different chromosomes, and so, while they co-regulate many genes, it is unlikely they were inherited as a pair. Knowing their interconnected roles in *V. harveyi* and *V. parahaemolyticus*, we wondered whether LuxT and SwrZ jointly regulate functions in other *Vibrionaceae* species. To garner preliminary evidence for or against this possibility, we examined the conservation of both genes within the *Vibrionaceae* family. Among the 418 sequenced *Vibrionaceae,* the *luxT* gene is present in all but 16 species (96%) (Fig 7A). None of the 16 species lacking *luxT* encodes a *swrZ* gene (Fig 7A). Among the 402 *Vibrionaceae* that possess *luxT,* 227 species (56%) also have *swrZ* (Fig 7A); these genera include *Aliivibrio, Photobacterium,* and *Vibrio* (Fig 7A). Thus, in cases in which both *luxT* and *swrZ* are present, the possibility for co-regulation of target genes exists. In the species that have *luxT* but lack *swrZ*, LuxT must regulate genes independently of SwrZ. Together, these observations may explain the evolution of SwrZ- dependent and SwrZ-independent functions for LuxT in *V. harveyi*.

**Fig 7.**
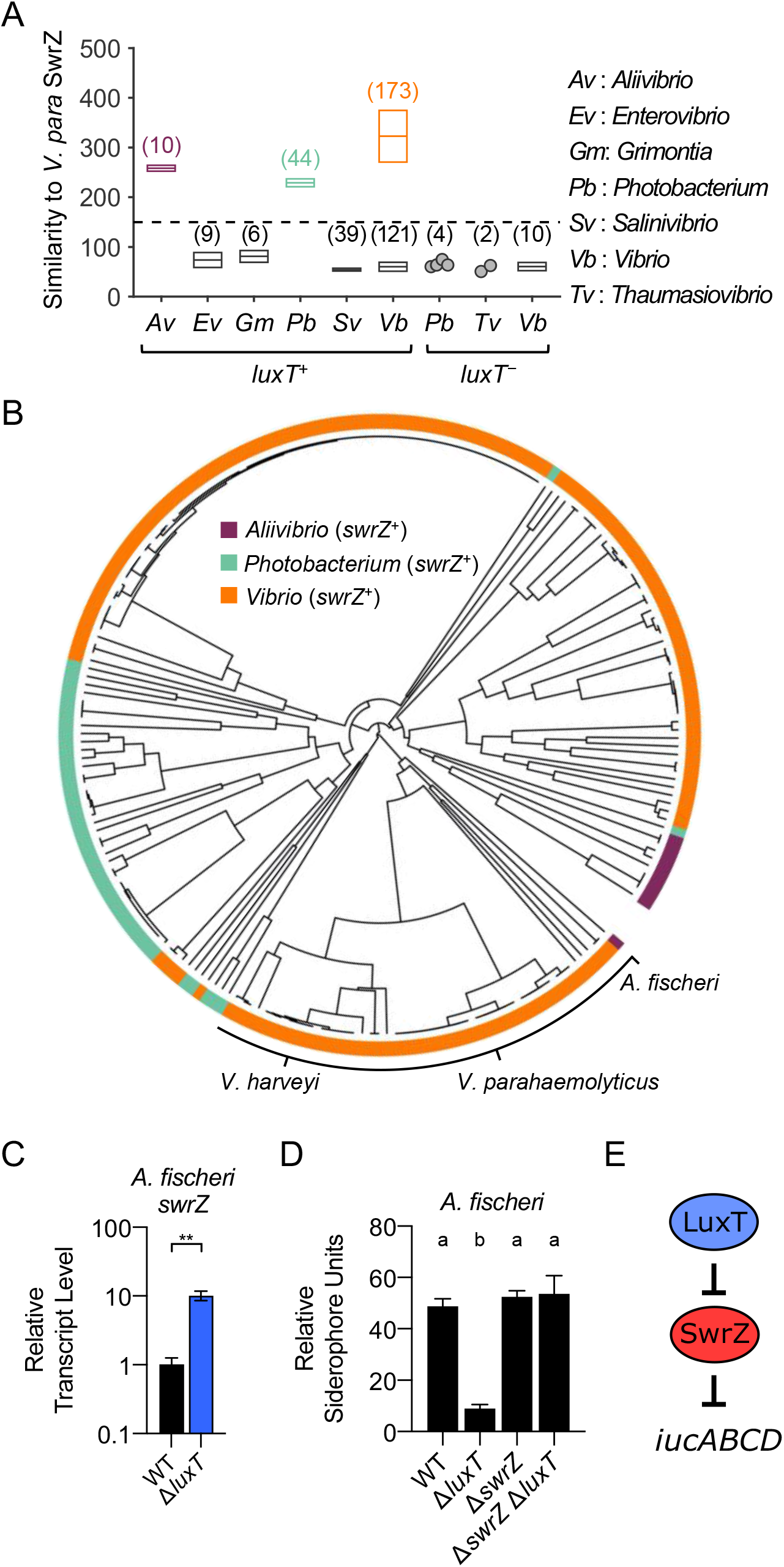
Co-occurrence of *luxT* and *swrZ* genes among *Vibrionaceae* species. **(A)** Highest similarity in protein sequences (BLOSUM62) in the indicated genera to *V. parahaemolyticus* SwrZ. Species are divided into two groups, one possessing *luxT* (*luxT*^+^) and one lacking *luxT* (*luxT*^−^). The dashed line indicates the cutoff used for the similarity score to differentiate *swrZ*^+^ from *swrZ*^-^ strains. Gray indicates groups of species that lack *swrZ*. The numbers in parentheses indicate the number of species in each group. Boxes show the means ± SDs for groups with > 5 species. Circles represent individual species in groups with ≤ 5 members. Purple, green, and orange denote, respectively, *Aliivibrio*, *Photobacterium*, and *Vibrio* species that possess *swrZ*. **(B)** Phylogenetic tree based on the *swrZ* promoter regions in *Vibrionaceae*. Phylogenetic distances were computed based on sequences spanning -122 to -8, relative to the *swrZ* start codons. Colors as in panel A. **(C)** qRT-PCR measurements of *swrZ* transcript levels in WT (black) and Δ*luxT* (blue) *A. fischeri.* RNA was isolated from strains grown in LM to OD_600_ = 0.1. **(D)** CAS assay quantitation of siderophore levels in cell-free culture fluids prepared from the indicated *A. fischeri* strains. Strains were grown for 16 h in AB medium. **(E)** Model for LuxT/SwrZ regulation of *A. fischeri* aerobactin siderophore production. In C and D, error bars represent standard deviations of the means of *n* = 3 biological replicates. In C, an unpaired two-tailed *t* test with Welch’s correction was performed comparing the indicated WT and Δ*luxT* conditions. *p*-value: ** < 0.01. In D, different letters indicate significant differences between strains, *p* < 0.05 (two-way ANOVA followed by Tukey’s multiple comparisons test).

To probe the possibility that LuxT also represses *swrZ* in other species, we performed a phylogenetic analysis by comparing the *swrZ* promoter regions of all *luxT^+^ swrZ*^+^ species. Our prediction was that *swrZ* promoters would be more similar in species in which LuxT represses *swrZ* than in species in which LuxT does not regulate *swrZ*. In support of this idea, the *V. harveyi* and *V. parahaemolyticus swrZ* promoter sequences are indeed similar. Thus, when we constructed a phylogenetic tree based on *swrZ* promoter sequences, *V. harveyi* and *V. parahaemolyticus* reside in close proximity (Fig 7B). As an initial test of LuxT regulation of *swrZ* in a species beyond *V. harveyi* and *V. parahaemolyticus*, we analyzed LuxT repression of *swrZ* in *Aliivibrio fischeri,* a species that neighbors *V. harveyi* and *V. parahaemolyticus* on the *swrZ* promoter phylogenetic tree (Fig 7B). Indeed, *swrZ* transcript levels were ∼10-fold higher in Δ*luxT A. fischeri* than in WT showing that LuxT is a repressor of *swrZ* in *A. fischeri* (Fig 7C). We previously reported that LuxT is an indirect activator of *iucABCD,* encoding the aerobactin siderophore biosynthesis genes in *A. fischeri* (21). At that time, we did not know the identity of the regulator connecting LuxT to *iucABCD*. Our findings here point to SwrZ fulfilling that function. To test this idea, we used the CAS assay to measure siderophore levels in culture fluids from WT, Δ*luxT,* Δ*swrZ,* and Δ*swrZ* Δ*luxT A. fischeri* strains. Indeed, LuxT activates *A. fischeri* siderophore production only in the *swrZ*^+^ background as deletion of *luxT* in the Δ*swrZ* strain caused no change in siderophore production (Fig 7D). Thus, LuxT activates *A. fischeri* siderophore production via a SwrZ-dependent mechanism. There is no evidence that a second SwrZ-independent mechanism occurs (Fig 7D). Our model for LuxT and SwrZ regulation of *A. fischeri* siderophore production is depicted in Fig 7E.

## Discussion

QS controls over 600 genes in *V. harveyi* (8). Integral to the *V. harveyi* QS process are five, largely redundant, Qrr sRNAs that post-transcriptionally regulate target gene expression. LuxT is a TetR family transcriptional regulator that was previously identified to repress expression of *V. harveyi qrr*1 encoding the Qrr1 sRNA (18). When LuxT acts as a *qrr*1 repressor, it tunes the expression of select Qrr1-controlled genes that are members of the much larger QS regulon. Here, we use RNA-Seq to discover that beyond controlling *qrr*1 expression, LuxT is a global regulator in *V. harveyi* that controls 414 genes. Analogous to what has been reported for *V. parahaemolyticus* (22), LuxT directly represses *swrZ,* encoding an additional transcriptional regulator. We show that, in *V. harveyi*, LuxT activates type III secretion, siderophore production, and aerolysin production genes independently of Qrr1, by both SwrZ-dependent and SwrZ-independent mechanisms. In the future, defining the SwrZ regulon will be a crucial step to gain a comprehensive understanding of the individual and combined roles of LuxT and SwrZ in regulating *V. harveyi* gene expression.

Two feedback loops exist in the *V. harveyi* LuxT/SwrZ regulatory pathway: LuxT and SwrZ activate and repress, respectively, their own transcription. Generally, positive feedback is thought to amplify responses to a stimuli, whereas negative feedback buffers pathway output against fluctuations in stimuli (45, 46). Thus, including both positive and negative feedback in a single system can alter both input sensitivity and output dynamics (46, 47). We speculate that the LuxT positive feedback loop enables *V. harveyi* to rapidly commit to the LuxT “ON” state following initial activation of *luxT* expression. By contrast, SwrZ-directed negative feedback on *swrZ,* which occurs when LuxT is in the “OFF” state, should prevent runaway *swrZ* expression. Moreover, LuxT and SwrZ have opposing regulatory functions, so genes activated by LuxT are repressed by SwrZ. Negative SwrZ autoregulation could, by tamping down SwrZ production, accelerate the expression of LuxT-activated genes when *V. harveyi* transitions to the LuxT “ON” state.

Our analyses revealed that LuxT activates T3SS, siderophore, and aerolysin genes by two mechanisms, one SwrZ-dependent and one SwrZ-independent. We propose two advantages to this regulatory arrangement. First, this dual mechanism may promote a “switch-like” response. Because SwrZ represses genes that are activated by LuxT, when LuxT is in the “ON” state, stronger target gene activation will occur if *swrZ* is repressed by LuxT than if not. Second, if expression and/or activity of either or both LuxT and SwrZ respond to additional regulatory inputs, linking the two nodes could promote more nuanced control of gene expression than if LuxT and *swrZ* were not connected in the circuit.

For *V. harveyi* to transition between LuxT “ON” and LuxT “OFF” states, a cue(s) must activate *luxT* expression or increase LuxT activity. We do not know the identity of this putative cue. We do know that LuxT is not regulated by QS and, consistent with this finding, *luxT* expression remains steady over the *V. harveyi* growth curve (Fig S3A and S3B). Curiously, however, genes in the LuxT-controlled regulon overlap with those in the QS-controlled regulon. In particular, LuxT activates expression of type III secretion, siderophore production, and aerolysin production genes, all of which are also QS-controlled and expressed at LCD, which is the typical pattern for QS-controlled virulence genes in vibrios (48). The above mentioned QS- and LuxT-regulated traits are all virulence factors that contribute to the success of vibrios as pathogens in host organisms (43, 49, 50). For example, in shrimp larvae, *V. harveyi* T3SS gene expression is 1,000- fold higher than that in laboratory culture (49). Possibly, *luxT* expression or LuxT activity is modulated by a host factor. Alternatively or in addition, TetR family members frequently bind small-molecule ligands (51). It could be that a *V. harveyi-,* host-, or environmentally-derived ligand promotes heightened LuxT activity. Future studies investigating regulation of *luxT* expression or LuxT activity should define the connection between the LCD state and LuxT function.

Our phylogenetic analyses revealed that *luxT,* but not *swrZ,* is conserved among *Vibrionaceae.* And, LuxT repression of *swrZ* is conserved at least in *V. parahaemolyticus, V. harveyi*, and our *A. fischeri* test case (22). Going forward, it will be interesting to learn how LuxT function has diverged in species that harbor *luxT* but lack *swrZ,* such as in *Vibrio cholerae* C6706. RNA-Seq experiments similar to those described here could be used to begin to parse the roles LuxT plays and, based on the findings, paint a coherent evolutionary picture of LuxT control of *Vibrionaceae* biology.

## Materials and Methods

### Bacterial strains, culture conditions, and standard methods

All strains are listed in Table S1A. *V. harveyi* strains were derived from *V. harveyi* BB120 (BAA-1116) (52). Previously, *V. harveyi* BB120 was reclassified as *Vibrio campbellii* BB120 (53). For literary consistency, we refer to this strain as *V. harveyi. A. fischeri* strains were derived from *A. fischeri* ES114 (54). *E. coli* S17–1 λ*pir* was used for cloning. *E. coli* MG1655 was used for heterologous gene expression. *V. harveyi* and *A. fischeri* strains were grown in either Luria Marine (LM) medium or minimal Autoinducer Bioassay (AB) medium at 30°C with shaking (55, 56). AB medium contained 0.4% vitamin-free casamino acids (Difco). *E. coli* strains were grown in LB medium at 37°C for cloning or at 30°C for heterologous gene expression. When necessary, kanamycin, chloramphenicol, ampicillin, and polymyxin B were added at 100 μg mL^-1^, 10 μg mL^-1^, 100 μg mL^-1^, and 50 μg mL^-1^, respectively. Gene expression from P*_BAD_* and P*_tac_* promoters was induced following addition of 0.01% arabinose and 0.5 mM isopropyl β-D-1-thiogalactopyranoside (IPTG), respectively, at the time of inoculation. Procedures for LuxT-6xHis purification, EMSA analyses, qRT-PCR measurements, and hemolysis assays have been described (18), as have CAS siderophore detection assays (21). Primers used to amplify DNA probes for EMSA analyses and for qRT-PCR measurements are listed in Table S1B. Growth conditions for qRT-PCR experiments are provided in the figure legends.

### DNA manipulation and strain construction

Oligonucleotide primers were purchased from Integrated DNA Technologies (IDT) and are listed in Table S1B. PCR reactions used either KOD Hot Start DNA Polymerase (Sigma) or iProof DNA Polymerase (Bio-Rad). All cloning was completed using Gibson Assembly Master Mix (New England Biolabs) for isothermal DNA assembly (57). Plasmids were verified by sequencing (Genewiz) and are listed in Table S1C. Regarding nomenclature of our constructs, a capital P designates the promoter driving transcription (e.g. P*_swrZ_*-*lux*). Plasmids that promote overexpression of genes are designated with a lowercase p (e.g. p*swrZ*). The P*_swrZ_*-*lux* and P*_luxT_*-*lux* transcriptional reporters included 114 and 531 bp promoter regions, respectively, to drive transcription of *luxCDABEG.* A consensus ribosome-binding site was included in both reporters to drive translation. Plasmids were introduced into *E. coli* by electroporation using a Bio-Rad Micro Pulser. Plasmids were conjugated into *V. harveyi* and *A. fischeri* from *E. coli* S17–1 λ*pir*. *V. harveyi* exconjugants were selected on agar plates containing polymyxin B, and *A. fischeri* exconjugants were selected on agar plates containing ampicillin. Chromosomal alterations in *V. harveyi* and *A. fischeri* were introduced using the pRE112 vector harboring the *sacB* counter-selectable marker as previously described (18, 58, 59). Mutations were verified by PCR and/or sequencing.

### RNA-Sequencing

Cells from overnight cultures of WT and Δ*luxT V. harveyi* strains grown in LM medium were pelleted by centrifugation at 21,100 x *g* (Eppendorf 5424) and resuspended in fresh LM medium. Fresh cultures containing 25 mL of LM medium were inoculated with the washed cells at OD_600_ = 0.005. The cultures were incubated at 30°C with shaking, and RNA was isolated from 3 biological replicates of each strain when the cultures had reached OD_600_ = 0.1 using the RNeasy mini kit (Qiagen #74106). RNA-Seq was performed at the Genomics Core Facility at Princeton University, as previously described (60). Reads were mapped to the *V. harveyi* BB120 (BAA-1116) genome using TopHat (61). Genes with differential expression were identified using DESeq2 (62) and those exhibiting a log_2_ fold-change in expression > 1, along with a *p*-value < 0.01, in the Δ*luxT* strain compared to WT *V. harveyi,* were designated as members of the LuxT regulon.

### Bioluminescence assays

Cells in overnight cultures of *E. coli* harboring the the P*_swrZ_*-*lux* reporter or *V. harveyi* harboring the P*_luxT_*-*lux* reporter were pelleted by centrifugation at 21,100 x *g* (Eppendorf 5424) and resuspended in LB or LM medium, respectively. The *E. coli* and *V. harveyi* cells were inoculated into fresh LB and LM, respectively, with normalization to a starting OD_600_ of 0.005. 150 μL of each culture was transferred to wells of a 96-well plate (Corning) in quadruplicate technical replicates and overlayed with 50 μL of mineral oil (Sigma). The plates were incubated with shaking at 30°C, and bioluminescence and OD_600_ were measured every 15 min for 24 h using a BioTek Synergy Neo2 Multi-Mode Reader (BioTek, Winooski, VT, USA). Relative light units (RLU: bioluminescence/OD_600_) represent the values when each sample was at OD_600_ = 1.

### VopD Western blot analyses

Cytoplasmic VopD levels were detected in *V. harveyi* cultures that had been grown in AB medium supplemented with 5 mM EGTA as previously described (5). Cells equivalent to 1 OD_600_ were pelleted by centrifugation at 21,100 x *g* (Eppendorf 5424), resuspended in 100 μL of SDS-PAGE sample buffer (Bio-Rad), and boiled for 10 min. Samples were loaded onto 4-20% Mini-PROTEAN TGX Gels (Bio-Rad) and subjected to electrophoresis for 30 min at 50 mA. Following Western transfer, nitrocellulose membranes were cut in half. One portion was probed with an antibody against VopD, and the other portion was probed with an antibody against the LuxS control, as previously described (5). Anti-rabbit IgG (H+L) HRP Conjugate (Promega) was used as the secondary antibody. Proteins were visualized using SuperSignal West Femto Maximum Sensitivity Substrate (Thermo Fisher) and an ImageQuant LAS 4000 imager.

### Phylogenetic analyses

Genomic DNA sequences of 418 *Vibrionaceae* species were downloaded from the GenBank database (63). Genes encoding *luxT* among *Vibrionaceae* species were identified previously (18). A custom MATLAB (Mathworks, 2020) search algorithm based on protein sequence similarity was used to identify genes encoding *swrZ*. Briefly, protein sequences of SwrZ from *V. harveyi* ATCC BAA-1116 and *V. parahaemolyticus* RIMD 2210633 were used as queries. Both yielded similar search results. The chromosomes or contigs of species under consideration were first converted to amino acid sequences, and subsequently scanned for regions similar to the query sequences. To identify regions of highest similarity between protein sequences, local sequence alignments were performed using the Smith-Waterman (SW) algorithm (64). The standard substitution matrix BLOSUM62 (https://ftp.ncbi.nih.gov/blast/matrices/) was used to compute similarity scores *S*, which considers both the length and sequence similarity of the alignment. A cut-off value *S* >150 was used to identify putative genes encoding SwrZ. Genes identified as *swrZ* homologs were verified to encode GntR-family transcriptional regulators. Species lacking *swrZ* were excluded from further phylogenetic analyses. The protocols used to perform phylogenetic analyses and tree building have been described previously (18).

### Data availability

All relevant data are within the manuscript and the Supplemental Information files. RNA-Sequencing data and files containing the numerical data for the main text and supplemental figures are provided at Zenodo (https://doi.org/10.5281/zenodo.5719716).

## Acknowledgments

RNA-Sequencing was performed at the Genomics Core Facility in the Lewis-Sigler Institute for Integrative Genomics at Princeton University. This work was supported by the Howard Hughes Medical Institute, National Institutes of Health (NIH) Grant 5R37GM065859, and National Science Foundation Grant MCB-1713731 (to BLB). MJE was supported by the NIH graduate training grant (NIGMS T32GM007388). The funders had no role in study design, data collection and interpretation, or the decision to submit the work for publication. The authors have no competing interests to declare. We thank members of the Bassler laboratory for insightful discussions and ideas. We lovingly acknowledge Jian-Ping Cong and her fundamental contributions to the work reported here. We dedicate this manuscript to her memory. We are grateful for her decades of devotion to advancing research in the bacterial cell-cell communication field and for her unflagging kindness to and support of former and current Bassler laboratory members.

## Main Figures

**Data Set S1. The *V. harveyi* LuxT regulon.** Genes activated (Sheet 1) and repressed (Sheet 2) by LuxT in *V. harveyi.* FC designates fold-change in gene expression in the Δ*luxT* strain compared to WT.

**Table S1.** Strains, primers, and plasmids used in this study. (A) Strain list, (B) primer list, and (C) plasmid list.

---

**Fig S1.**
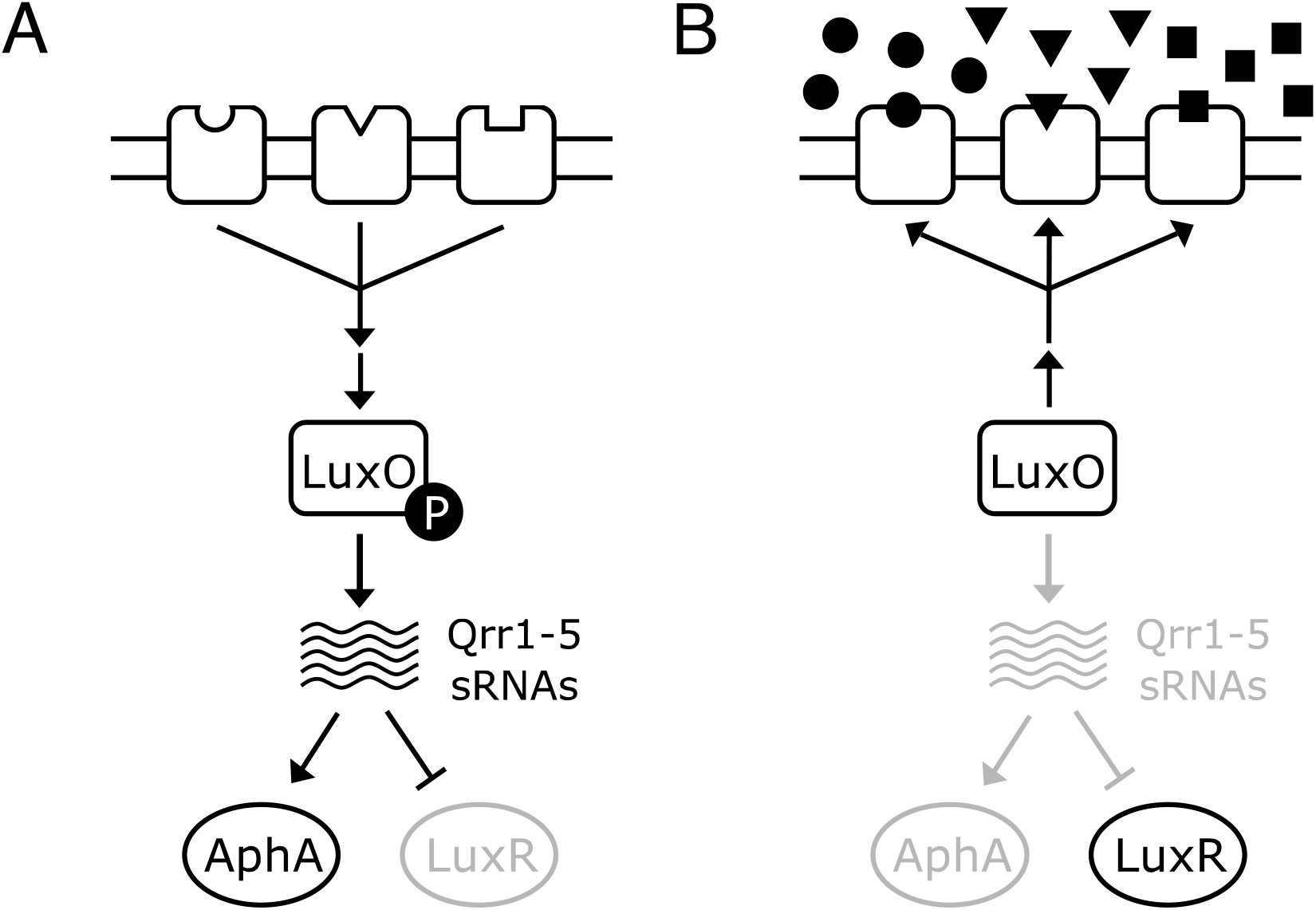
Simplified *V. harveyi* QS pathway. **(A)** LCD and **(B)** HCD *V. harveyi* QS circuit. Circles, triangles, and squares represent AIs. See Introduction in the main text for details.

**Fig S2.**
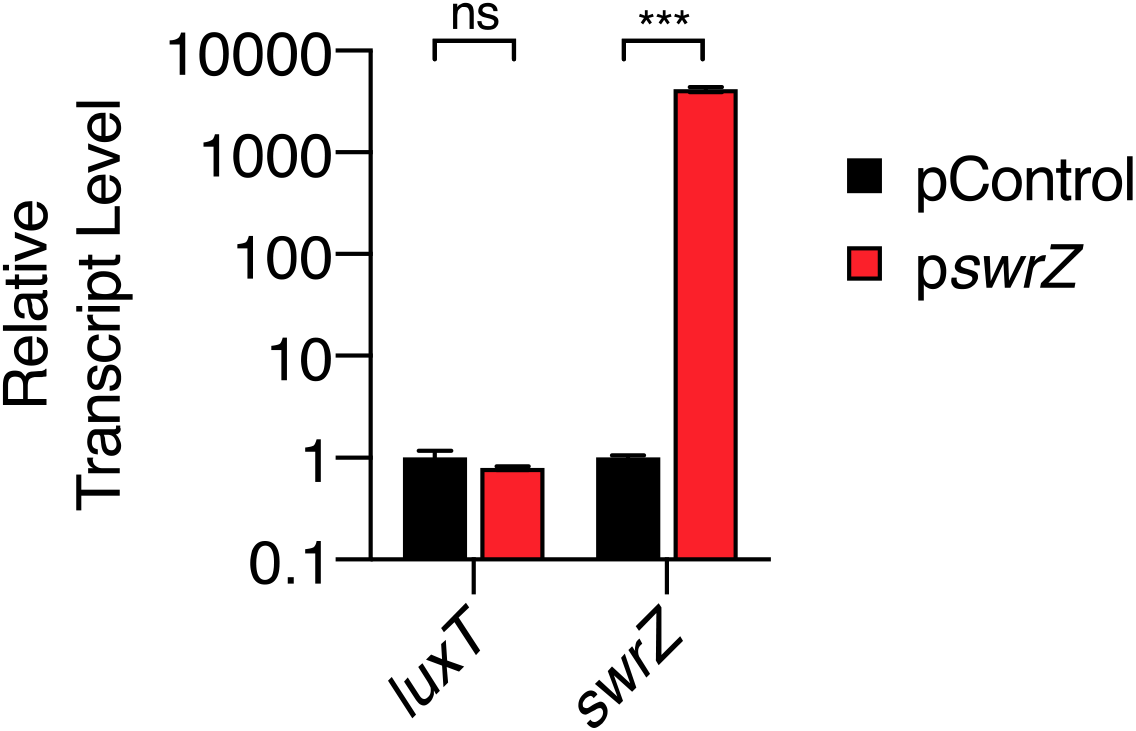
SwrZ does not regulate *luxT*. qRT-PCR measurements of *luxT* and *swrZ* transcript levels in WT *V. harveyi* harboring the following plasmids: pControl (black) denotes the empty parent vector and p*swrZ* (red) denotes a vector carrying IPTG-inducible *swrZ.* RNA was isolated from strains grown in LM containing 0.5 mM IPTG to OD_600_ = 1. Unpaired two-tailed *t* tests with Welch’s correction were performed comparing the pControl and p*swrZ* conditions for each gene. *p-*values: ns ≥ 0.05, *** < 0.001.

**Fig S3.**
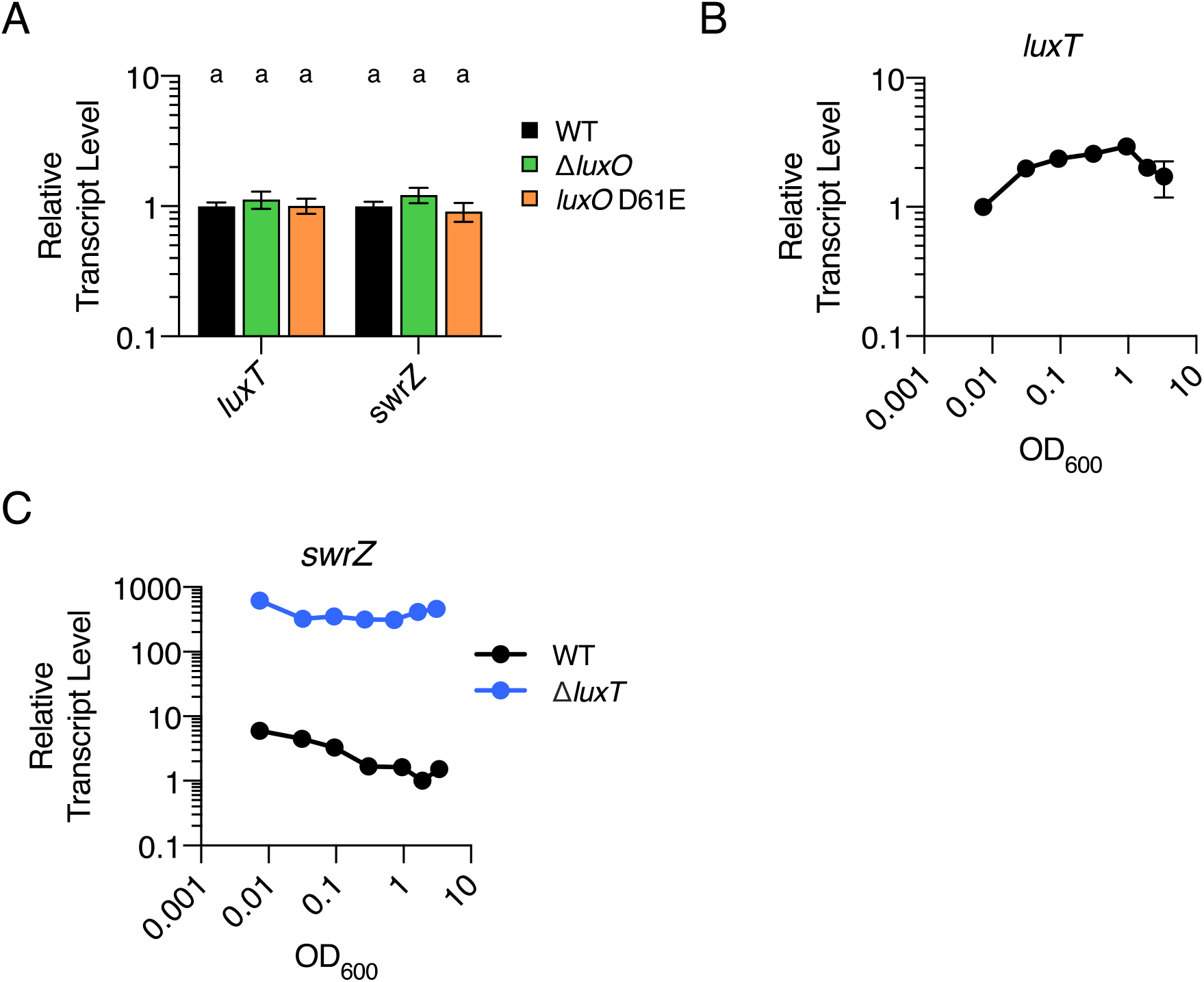
QS does not regulate *luxT* or *swrZ* in *V. harveyi.* **(A)** qRT-PCR measurements of *luxT* and *swrZ* transcript levels in WT (black), Δ*luxO* (green), and *luxO* D61E (orange) *V. harveyi* strains. RNA was isolated from strains grown in AB to OD_600_ = 1. **(B)** qRT-PCR measurements of *luxT* transcript levels over growth. RNA was isolated from WT *V. harveyi* grown in LM to different cell densities, as indicated by the OD_600_ values on the x-axis. **(C)** As in B, except *swrZ* transcript levels were measured in WT (black) and Δ*luxT* (blue) *V. harveyi* strains. In all panels, error bars represent standard deviations of the means of *n* = 3 biological replicates. Standard deviations that are smaller than the symbols are not shown. In A, different letters indicate significant differences between strains, *p* < 0.05 (two-way analysis of variation (ANOVA) followed by Tukey’s multiple comparisons test).

**Fig S4.**
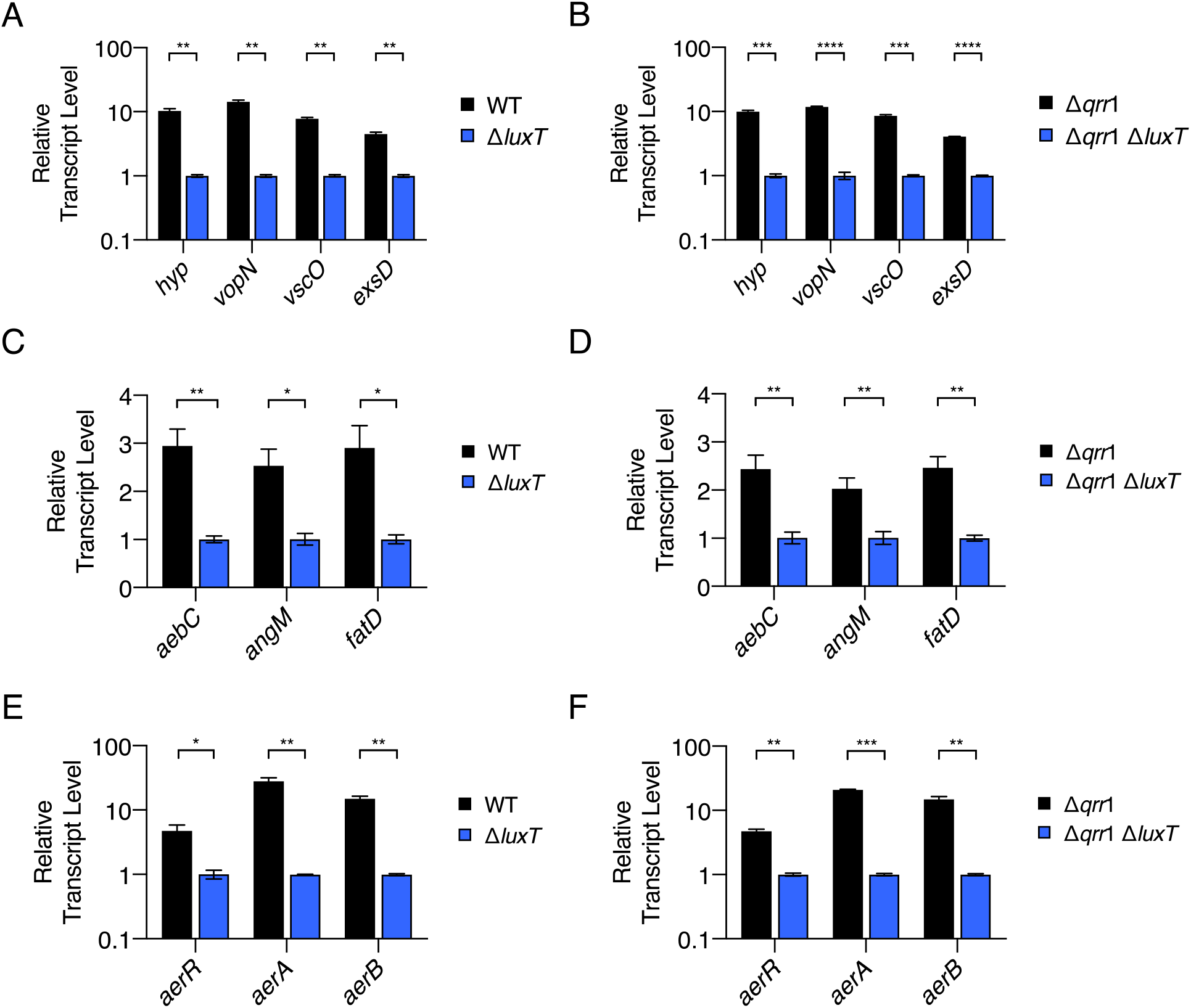
LuxT activates T3SS, siderophore, and aerolysin genes by Qrr1- independent mechanisms in *V. harveyi*. qRT-PCR measurements of transcript levels of the indicated genes in WT (black) and Δ*luxT* (blue) *V. harveyi* **(A, C,** and **E)** or in Δ*qrr*1 (black) and Δ*qrr*1 Δ*luxT* (blue) *V. harveyi* strains **(B, D,** and **F)**. RNA was isolated from strains grown in LM to OD_600_ = 0.1. In all panels, error bars represent standard deviations of the means of *n* = 3 biological replicates. Unpaired two-tailed *t* tests with Welch’s correction were performed comparing two conditions, as indicated. *p*-values: * <0.05, ** < 0.01, *** < 0.001, **** < 0.0001.

**Fig S5.**
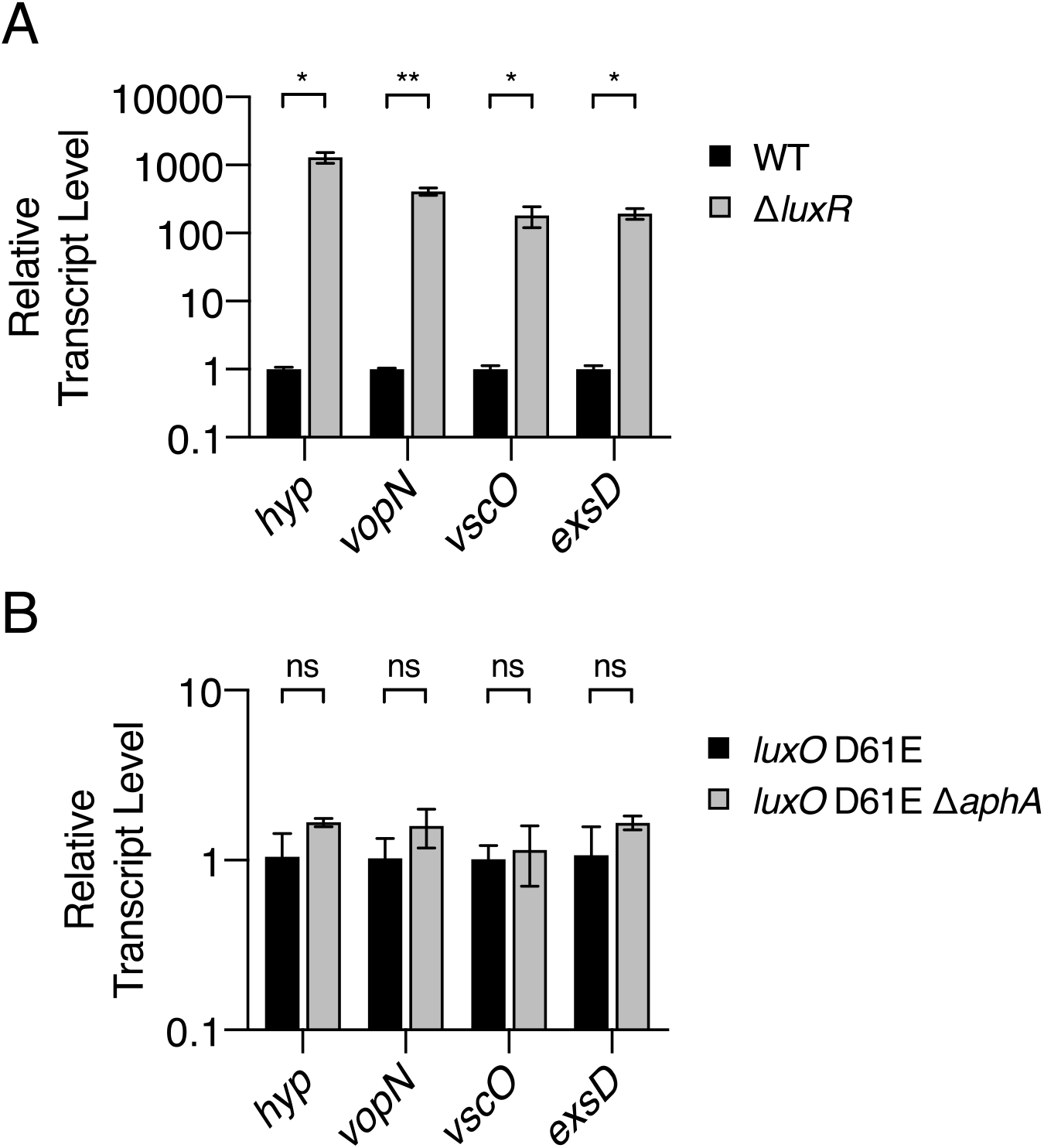
*V. harveyi* T3SS genes are repressed at HCD by LuxR. **(A)** qRT-PCR measurements of transcript levels of the indicated T3SS genes in WT (black) and Δ*luxR* (gray) *V. harveyi* strains. RNA was isolated from strains grown in AB to OD_600_ = 1. **(B)** As in A, except transcript levels were measured in *luxO* D61E (black) and *luxO* D61E Δ*aphA* (gray) *V. harveyi* strains. In both panels, error bars represent standard deviations of the means of *n* = 3 biological replicates. Unpaired two-tailed *t* tests with Welch’s correction were performed comparing two conditions, as indicated. *p*-values: ns ≥ 0.05, * < 0.05, ** < 0.01.

**Fig S6.**
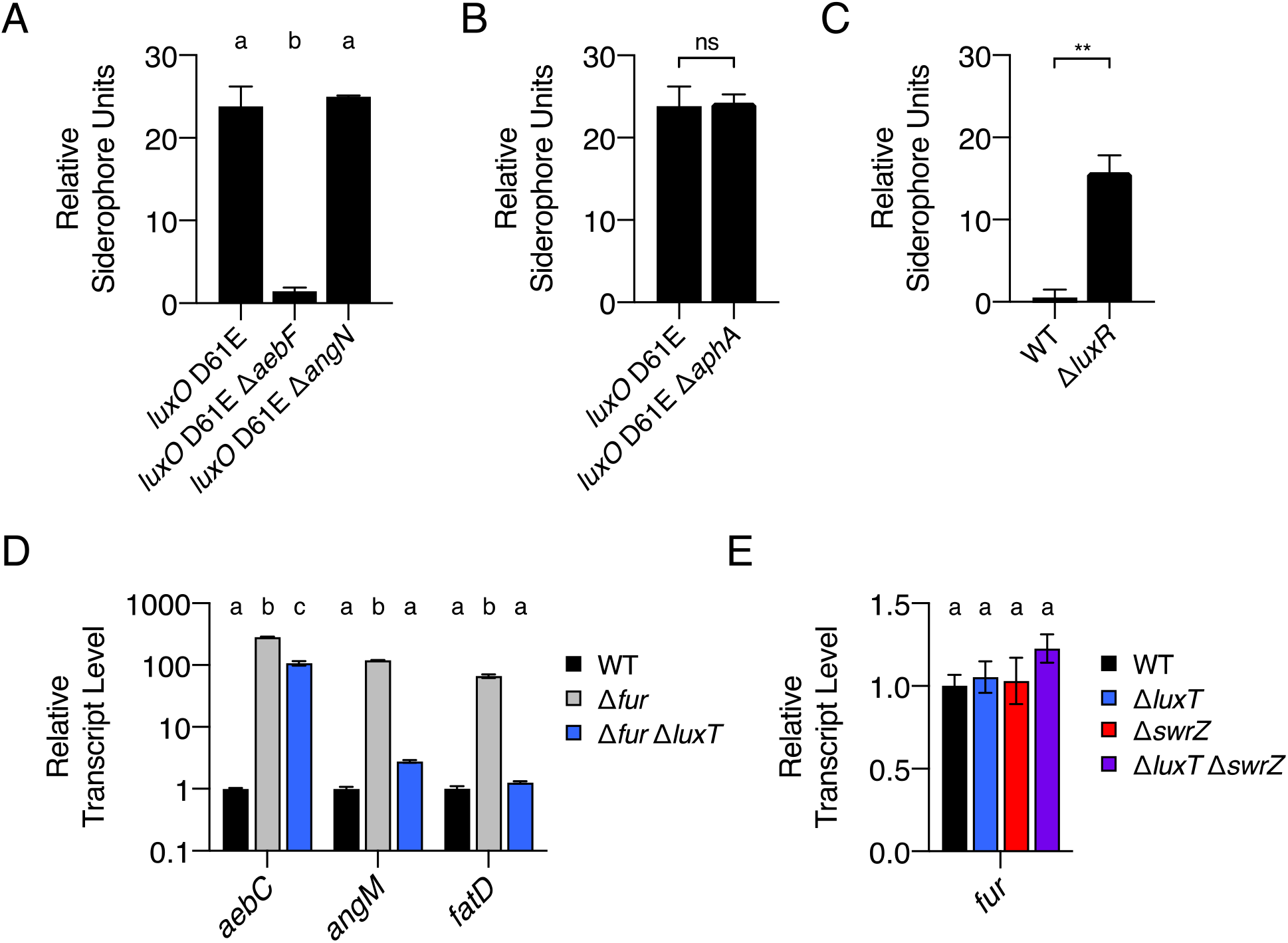
LuxR and Fur repress production of the *V. harveyi* amphi-enterobactin siderophore. **(A-C)** CAS assay quantitation of siderophore levels in cell-free culture fluids isolated from the indicated *V. harveyi* strains. Strains were grown for 16 h in AB. **(D)** qRT-PCR measurements of transcript levels of the indicated genes in WT (black), Δ*fur* (gray), and Δ*fur* Δ*luxT* (blue) *V. harveyi* strains. Strains were grown in LM to OD_600_ = 0.1. **(E)** As in D for WT (black), Δ*luxT* (blue), Δ*swrZ* (red), and Δ*swrZ* Δ*luxT* (purple) *V. harveyi* strains. In all panels, error bars represent standard deviations of the means of *n* = 3 biological replicates. In A, D, and E, different letters indicate significant differences between strains, *p* < 0.05 (two-way ANOVA followed by Tukey’s multiple comparisons test). In B and C, unpaired two-tailed *t* tests with Welch’s correction were performed comparing the two conditions, as indicated. *p*-values: ns ≥ 0.05, ** < 0.01.

**Fig S7.**
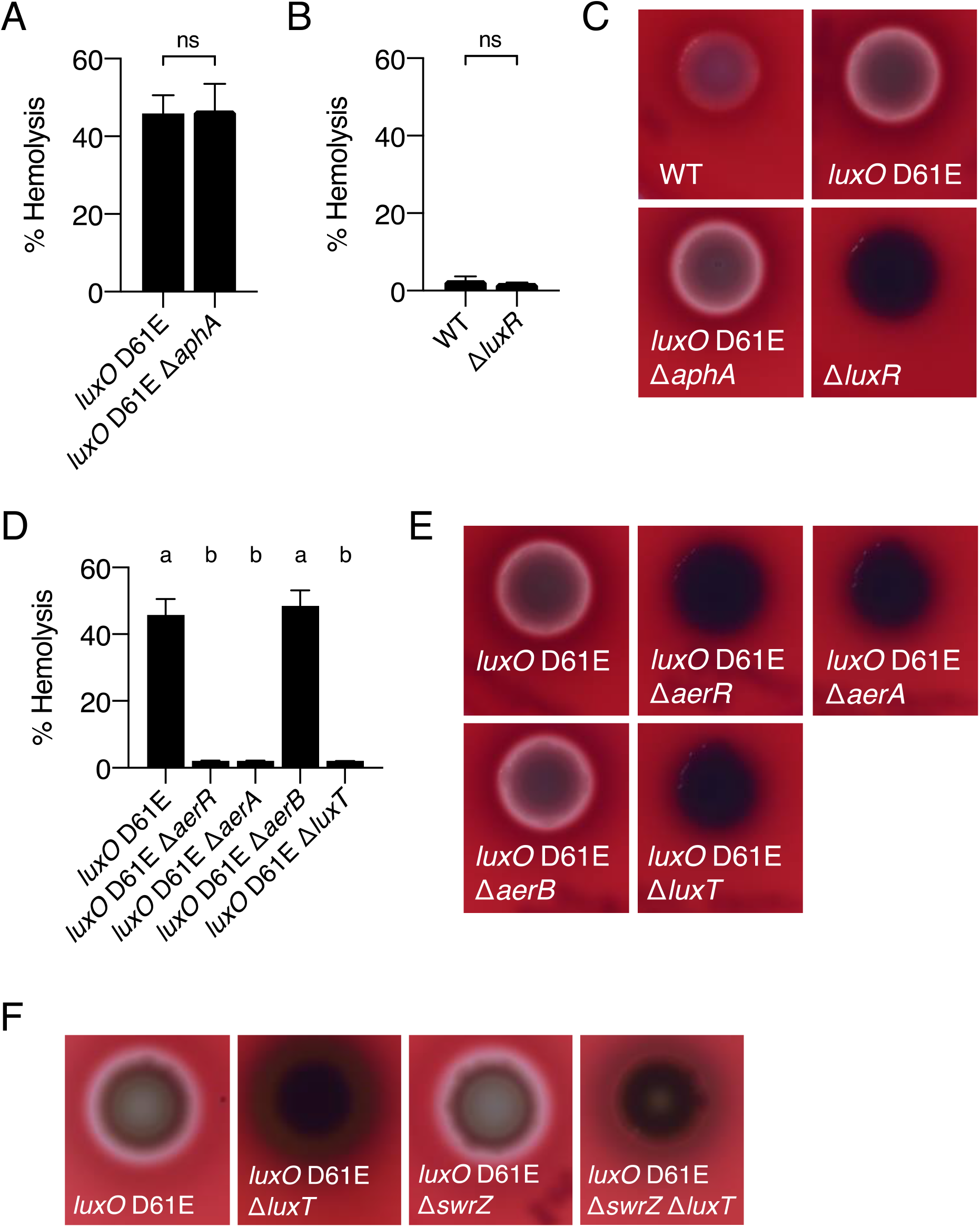
Assessment of the requirements for *V. harveyi* aerolysin production. **(A, B, and D)** Hemolytic activity present in the indicated *V*. *harveyi* cell-free culture fluids as judged by lysis of defibrinated sheep’s blood cells. Culture fluids were collected after 24 h of growth in AB medium. **(C, E,** and **F)** Hemolytic activity produced by the indicated *V. harveyi* strains as judged by halo formation on TSA plates containing 5% sheep’s blood. Strains were grown for 72 h at 30°C. In A, B, and D, error bars represent standard deviations of the means of *n* = 3 biological replicates. In A and B, unpaired two-tailed *t* tests with Welch’s correction were performed comparing two conditions, as indicated. *p*- values: ns ≥ 0.05. In D, different letters indicate significant differences between strains, *p* < 0.05 (two-way ANOVA followed by Tukey’s multiple comparisons test).

**Fig S8.**
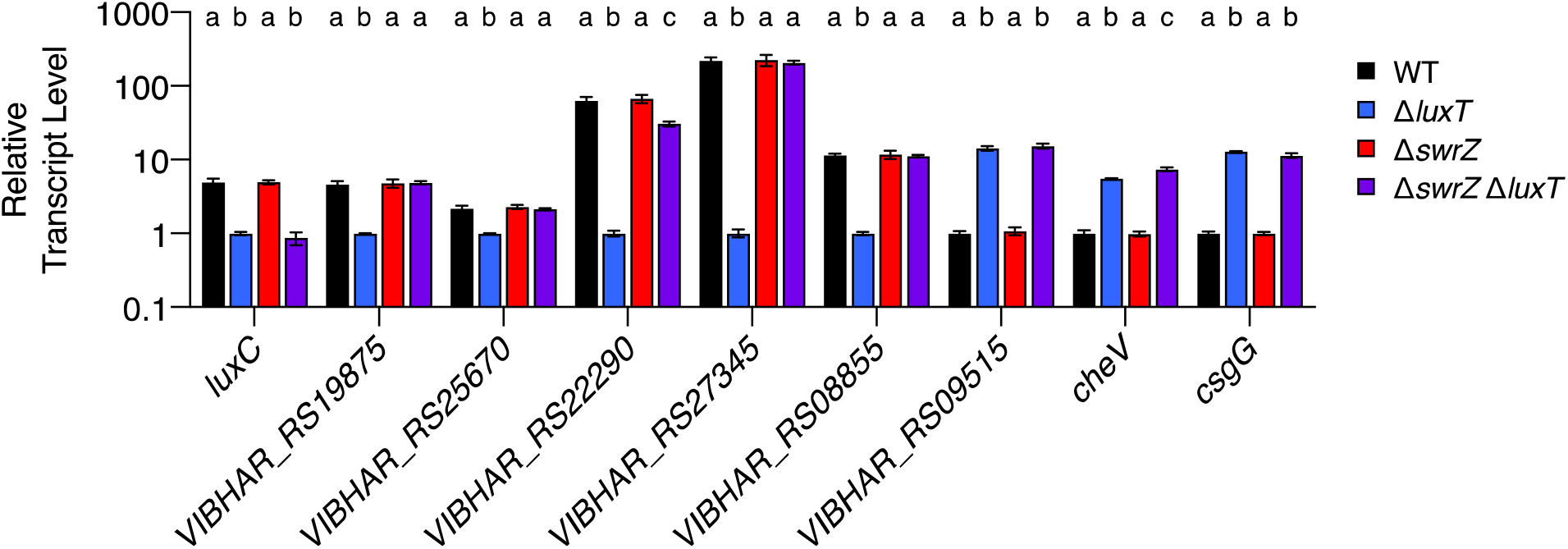
LuxT regulates genes by SwrZ-dependent and SwrZ-independent mechanisms in *V. harveyi.* qRT-PCR measurements of transcript levels of the indicated genes in WT (black), Δ*luxT* (blue), Δ*swrZ* (red), and Δ*swrZ* Δ*luxT* (purple) *V. harveyi* strains. Strains were grown in LM to OD_600_ = 0.1. Error bars represent standard deviations of the means of *n* = 3 biological replicates. Different letters indicate significant differences between strains, *p* < 0.05 (two-way ANOVA followed by Tukey’s multiple comparisons test).

